# A gut-restricted lithocholic acid analog as an inhibitor of gut bacterial bile salt hydrolases

**DOI:** 10.1101/2021.03.15.435552

**Authors:** Arijit A. Adhikari, Deepti Ramachandran, Snehal N. Chaudhari, Chelsea E. Powell, Megan D. McCurry, Alexander S. Banks, A. Sloan Devlin

## Abstract

Bile acids play crucial roles in host physiology by acting as both detergents that aid in digestion and as signaling molecules that bind to host receptors. Gut bacterial bile salt hydrolase (BSH) enzymes perform the gateway reaction leading to the conversion of host-produced primary bile acids into bacterially modified secondary bile acids. Small molecule probes that target BSHs will help elucidate the causal roles of these metabolites in host physiology. We previously reported the development of a covalent BSH inhibitor with low gut permeability. Here, we build on our previous findings and describe the development of a second-generation gut-restricted BSH inhibitor with enhanced potency, reduced off-target effects, and durable in vivo efficacy. SAR studies focused on the bile acid core identified a compound, **AAA-10**, containing a C3-sulfonated lithocholic acid scaffold and an alpha-fluoromethyl ketone warhead as a potent pan-BSH inhibitor. This compound inhibits BSH activity in conventional mouse fecal slurries, bacterial cultures, and purified BSH proteins and displays reduced toxicity against mammalian cells compared to first generation compounds. Oral administration of **AAA-10** to wild-type mice for 5 days resulted in a decrease in the abundance of the secondary bile acids deoxycholic acid (DCA) and lithocholic acid (LCA) in the mouse GI tract with low systemic exposure of **AAA-10**, demonstrating that **AAA-10** is an effective tool for inhibiting BSH activity and modulating bile acid pool composition in vivo.

---

Metabolites derived from the human gut microbiota have been implicated as causal agents in the maintenance of host health and the progression of disease.^1^ Advances in metabolomics, sequencing technologies, and the development of genetic tools have facilitated the identification of bacterial metabolites and the biosynthetic pathways responsible for their production. However, the lack of specific tools to control the levels of these metabolites in complex microbial communities has hindered our ability to interrogate the roles of these metabolites in host physiology. Encouragingly, the recent development of small molecule modulators of bacterial metabolites has revealed the potential of microbiota-targeted therapies to treat disease, including colon cancer, cardiovascular disease, and Parkinson’s disease.^2-4^

Bile acids are one large class of molecules that undergo substantial metabolism by gut bacteria. ^5^ While bile acids have long been studied for their detergent properties,^6,7^ recent findings have illustrated the key role that these metabolites play as signaling molecules. Specific bile acids act as ligands for host nuclear hormone receptors (NhRs) and G-protein-coupled receptors (GPCRs), thereby affecting host metabolic and immunomodulatory processes.^8-11^ Disruption of bile acid homeostasis has been implicated in the initiation and progression of disease, including cancer, obesity, and hypercholesterolemia,^8,12-15^ underscoring the need for tools that control the levels of these metabolites in vivo.

Host-produced primary bile acids are conjugated to taurine or glycine in the mammalian liver, stored in the gallbladder, and secreted into the small intestine post-prandially where they act as detergents that facilitate digestion. In the lower GI tract, resident bacteria chemically modify these metabolites, producing a large class of molecules called secondary bile acids. Before these modifications can occur, the C24 amide of conjugated bile acids must be hydrolyzed, a gateway reaction that is carried out exclusively by gut bacterial bile salt hydrolases (BSHs) (EC 3.5.1.24)(Figure 1A).^16^ BSHs are widespread in human gut bacteria and have been identified in members of 12 different phyla, including Bacteroidetes and Firmicutes, the two dominant phyla in the human gut.^17^ Recent studies have found that BSH abundance or activity are correlated with human diseases, including inflammatory bowel diseases, type 2 diabetes, and cardiovascular disease.^17-19^ The causal role of BSH activity in host physiology, however, remains unclear. For example, studies involving antibiotic-treated and germ-free mice colonized with BSH-containing or BSH-deficient bacteria^20,21^ or conventional mice treated with non-selective small molecules^22,23^ have reported conflicting results about the effects of BSH activity on host metabolism. An inhibitor that targets a wide array of BSHs but exhibits limited off-target effects against bacterial and host cells would allow for the selective in vivo modulation of bile acid composition, shifting the bile acid pool toward conjugated bile acids and decreasing the abundance of deconjugated and secondary bile acids. Such a tool could be utilized in fully colonized animals and would provide valuable information about how bile acids affect host physiology.

**Figure 1.**
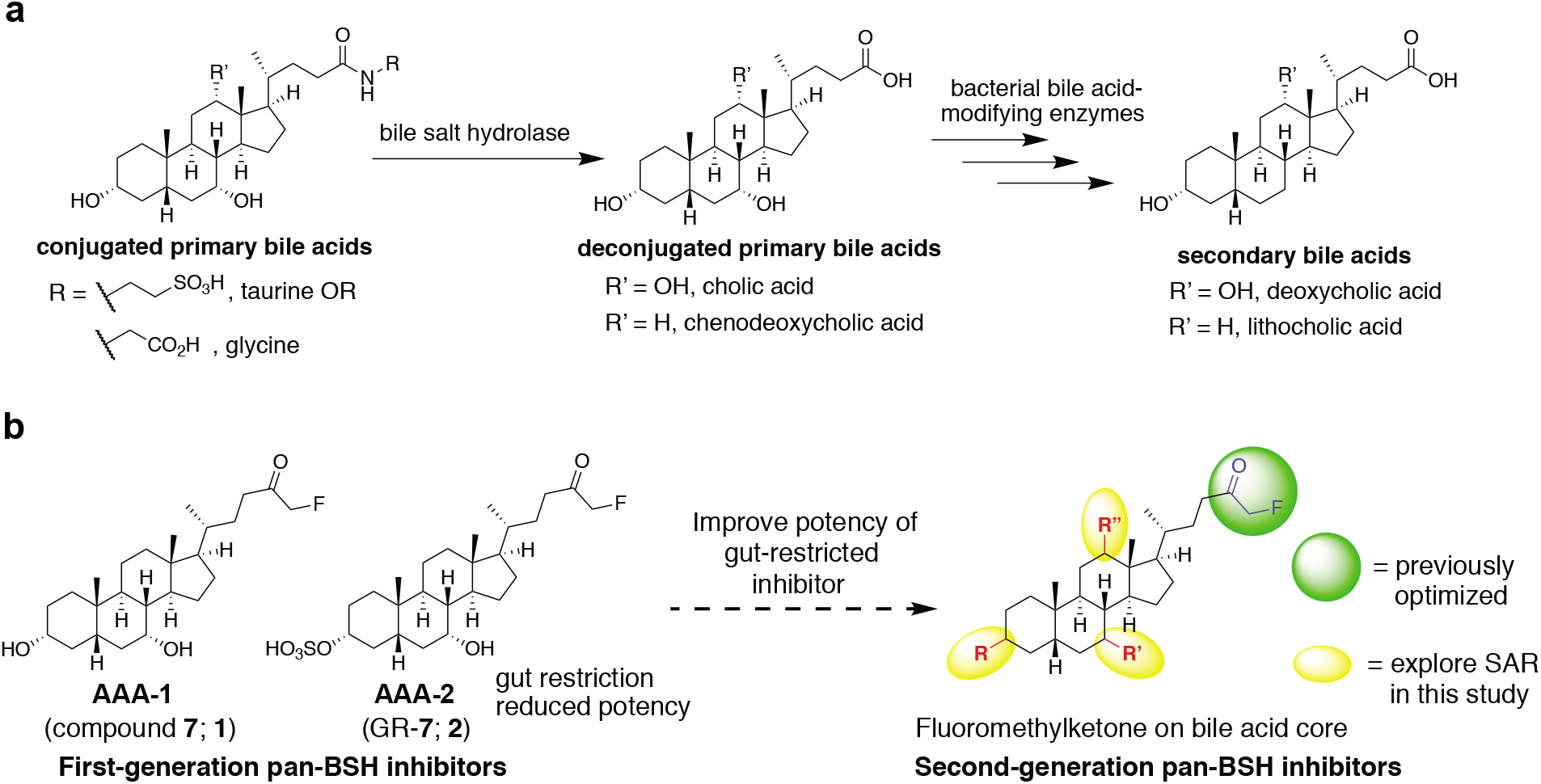
Targeting gut bacterial BSHs. **a**, Bacterial bile salt hydrolases (BSHs) perform the gateway reaction leading from host-produced conjugated primary bile acids to bacterially modified secondary bile acids. **b**, Development of second-generation BSH inhibitors starting from previously reported covalent pan-BSH inhibitors. Sulfonation of **AAA-1** at the C3-OH position previously resulted in an inhibitor with low systemic exposure but decreased potency (**AAA-2**). Here, SAR studies focusing on the bile acid core were performed with the goal of yielding a second-generation pan-BSH inhibitor with improved potency.

In prior work, we took advantage of the nucleophilicity of the highly conserved active site N-terminal cysteine residue (Cys2) in BSHs to develop first-in-class covalent pan-BSH inhibitors (Figure 1B).^24^ In this study, we screened electrophilic warheads appended to the core of chenodeoxycholic acid (CDCA), an abundant human bile acid that is recognized by a broad spectrum of BSHs.^20,25,26^ This work established an alpha-fluoromethylketone (FMK)-containing molecule, compound **7** (referred to here as **AAA-1, 1**), as a potent and selective pan-BSH inhibitor. Treatment of conventional mice with a single dose of **AAA-1** allowed us to inhibit BSH activity and shift the in vivo bile acid pool toward host-produced bile acids for one day. We also showed that appending a sulfonate^27^ to the C3 hydroxyl group resulted in gut-restriction of the inhibitor, a change that limited the systemic exposure of this compound (GR-**7**, referred to here as **AAA-2, 2**). These studies demonstrated the potential of alpha-fluoromethyl ketone-containing inhibitors to target BSHs in vivo.

To increase the utility of BSH inhibitors for use in vivo as well as to overcome several limitations in our prior study, we sought to develop second-generation inhibitors. The gut-restricted inhibitor **AAA-2** exhibited lower potency than **AAA-1**, motivating the synthesis of new lead compounds. Moreover, while we previously demonstrated proof-of-principle that a gut-restricted inhibitor could affect BSH activity in vivo, we did not demonstrate a shift in the in vivo bile acid pool in our prior work. Finally, to demonstrate the potential utility of these compounds in animal models, we sought to show that a gut-restricted inhibitor could shift the bile acid pool over a multi-day period. Here, we have built on our previous findings and report the development of a second-generation gut-restricted BSH inhibitor with enhanced potency, reduced off-target effects, and multi-day in vivo efficacy. Our structure-activity relationship (SAR) studies focused on the bile acid core, and we identified a lithocholic acid core-based inhibitor, **AAA-10** (**8**) (Figure 2), as a potent pan-BSH inhibitor through screening against conventional mouse fecal slurries, bacterial cultures, and recombinant proteins. This compound is not antibacterial, displays reduced toxicity against mammalian cells compared to **AAA-1** and **AAA-2**, and does not affect signaling through the farnesoid X receptor (FXR) or Takeda G-protein receptor 5 (TGR5), key bile acid-mediated receptors. Finally, we demonstrate that **AAA-10** (**8**) can modulate the in vivo bile acid pool for 5 days, resulting in the decreased abundance of the secondary bile acids deoxycholic acid (DCA) and lithocholic acid (LCA).

**Figure 2.**
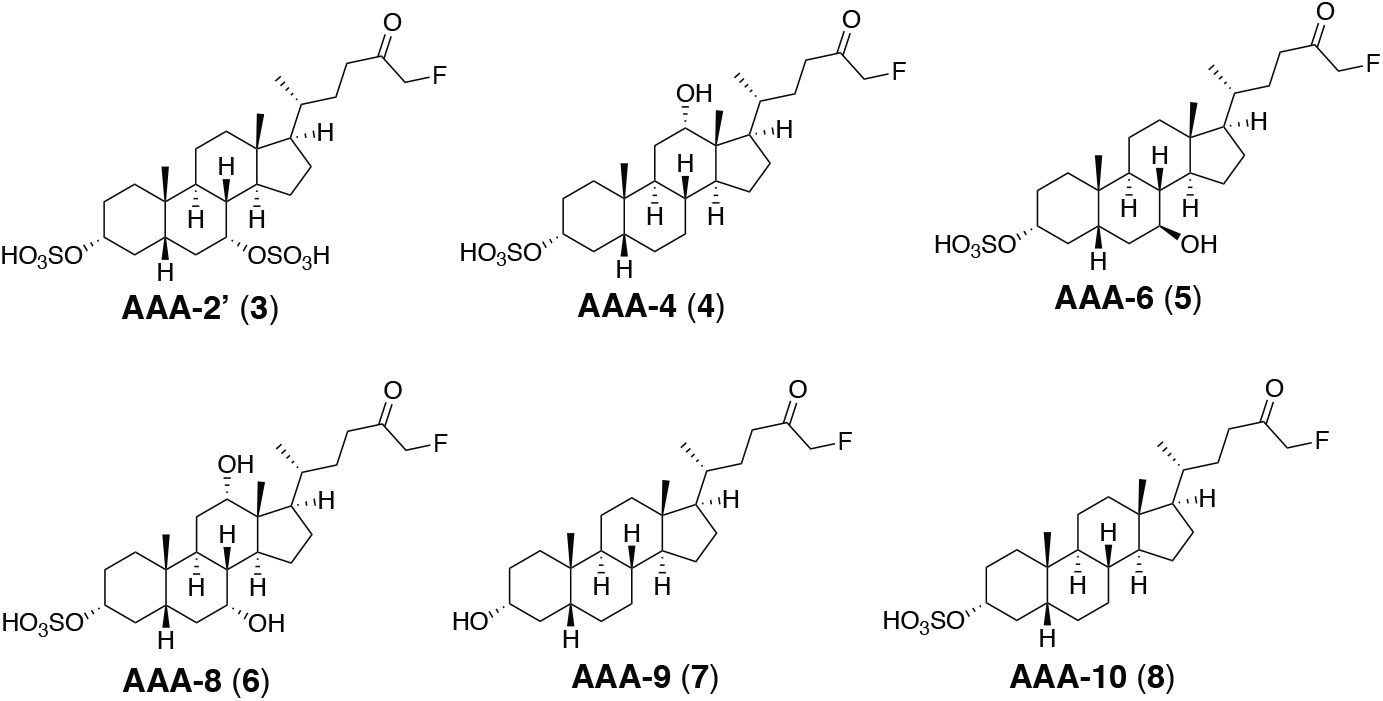
Library of sulfonated inhibitors. A small library of inhibitors was generated with SAR focused on incorporating the cores of naturally occurring bile acids found in both mouse and humans while maintaining an α-fluoromethyl ketone electrophile.

## RESULTS AND DISCUSSION

### Synthesis of BSH inhibitor candidates

In prior work, sulfonation of **AAA-1** (**1**) at C3, a position that is exposed to solvent in the co-crystal structure of this compound with the BSH from the gut bacterium *Bacteroides thetaiotaomicron* (*B. theta*), increased the solubility of this compound and limited its systemic exposure.^24,28^ The resultant compound **AAA-2**, (**2**), however, was less potent than **AAA-1**.^24^ With the goal of improving potency while still maintaining gut restriction, we decided to append the optimized FMK warhead on naturally occurring bile acid cores found in both the murine and human gut (CDCA, DCA, ursodeoxycholic acid (UDCA), cholic acid (CA), and LCA). These compounds could then be sulfonated at the C3 position to produce second-generation inhibitor candidates (Figure 1B).

In order to expedite the synthesis of the library, an optimized protecting group-free synthesis was developed (Scheme S1).^29^ Following activation of the unprotected bile acid with CDI, addition of magnesium benzyl fluoromalonate provided the fluoro beta-ketoester. Removal of the benzyl group followed by decarboxylation under hydrogenation conditions provided the FMK compounds. Finally, the sulfonate group was installed using SO_3_.pyridine. Because the C3-α-OH group on the bile acid core is more sterically accessible than the C7 and C12 α-alcohols, the sulfonation reactions proceeded selectively to provide the candidate C3-sulfonated inhibitors (Figure 2, **3-6, 8**).

### Library screen in conventional mouse feces

We previously reported the use of a wild-type conventional mouse fecal slurry assay^23^ to identify pan-inhibitors of BSHs.^24^ Because fecal slurry should contain BSHs from nearly all bacteria in the distal region of the murine GI tract, demonstrating inhibition of BSH activity in this assay represents an important benchmark that all inhibitor candidates should meet. We therefore utilized this assay as the first screen in the process of developing second-generation BSH inhibitors. Inhibitor candidates were added to fresh feces obtained from conventional wild-type mice (C57Bl/6J) and suspended in buffer under reducing conditions (Figure 3A). To facilitate identification of an inhibitor with enhanced potency compared to **AAA-2**, the first-generation gut-restricted inhibitor, compounds were intentionally tested at 10 µM, a concentration at which neither **AAA-1** nor **AAA-2** completely inhibits enzyme activity.^24^ After 30 min, glycine-conjugated deuterated chenodeoxycholic acid (GCDCA-*d*4) was added and its conversion to the deconjugated product CDCA-*d4* was quantified using ultra-high performance liquid chromatography-mass spectrometry (UPLC-MS).

**Figure 3.**
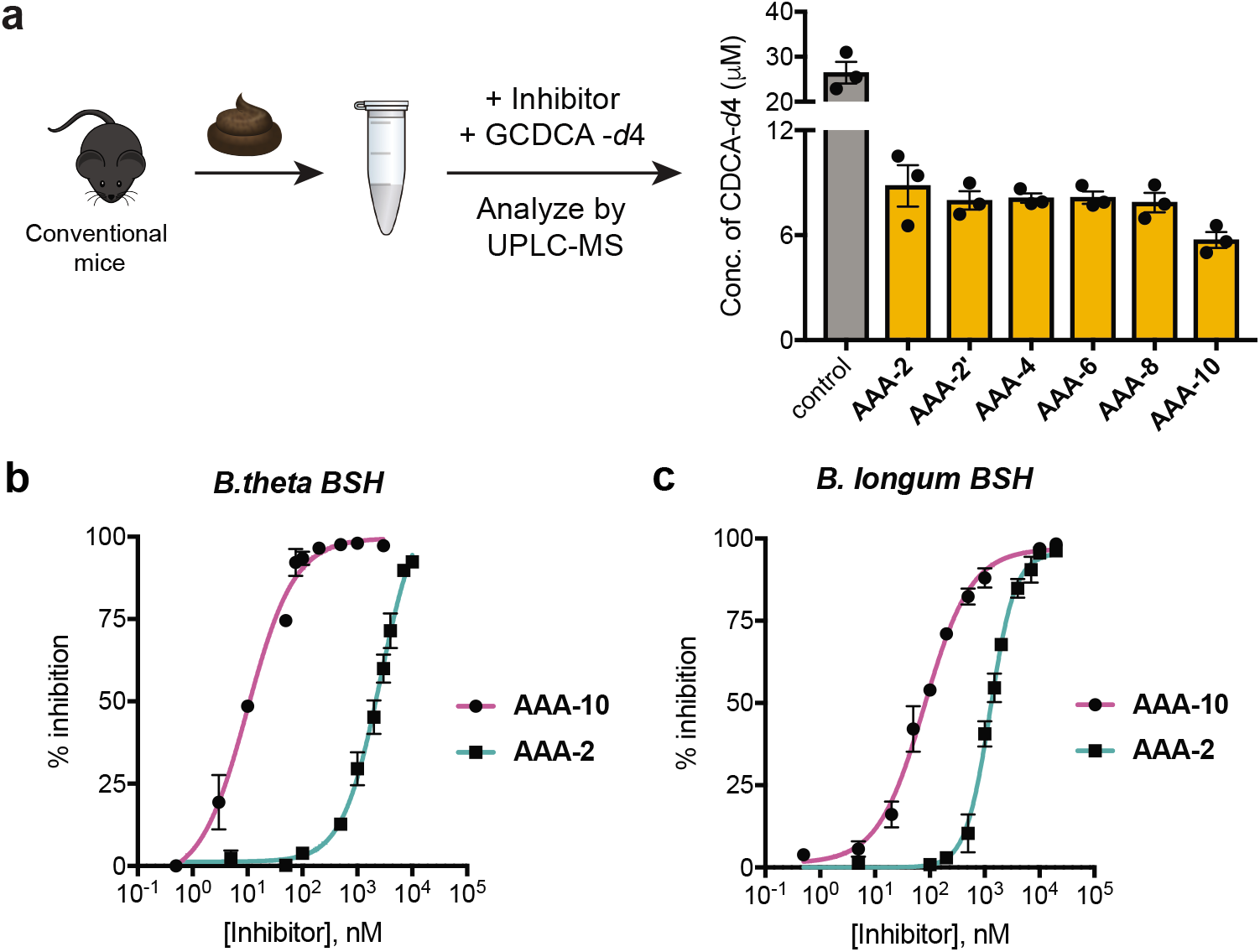
Identification of AAA-10 as a second-generation pan-BSH inhibitor. **a**, Assay design for screening the inhibitor library. Screening in fresh mouse feces identified **AAA-10** as a potent second-generation pan-inhibitor of BSHs. Inhibitors were tested at a concentration of 10 μM. Assays were performed three times independently in biological triplicate with similar results. **b** and **c, AAA-10** is more potent than **AAA-2** against recombinant BSHs. Comparison of **AAA-10** and **AAA-2** dose-response curves against *Bacteroides thetaiotaomicron (B. theta)* and *Bacteroides adolescentis (B. adolescentis)* BSHs using tauro-ursodeoxycholic acid (TUDCA) and tauro-deoxycholic acid (TDCA) as the respective substrates. See Table S1 for comparison of IC_50_ values of **AAA-1, AAA-2** and **AAA-10**. For **b** and **c**, Graphpad was used to fit IC_50_ curves. All assays were performed in biological triplicate, and data are presented as mean ± s.e.m.

Mono-sulfonated inhibitor candidates containing DCA, UDCA, and CA cores (**AAA-4** (**4**), **AAA-6** (**5**), and **AAA-8** (**6**), respectively) inhibited BSH activity but were not more potent than **AAA-2** in this assay (Figure 3A). The C3, C7-disulfonated derivative **AAA-2’** (**3**) was also equipotent to **AAA-2**. Notably, the LCA core-based analog **AAA-10** was more potent than both **AAA-2** and the inhibitor candidates **AAA-2’, AAA-4, AAA-6**, and **AAA-8. AAA-10** was equipotent to its unsulfonated analog **AAA-9** (**7**), indicating that C3-sulfonation did not hinder BSH inhibitory activity (Figure S1A).

Mice fed a high-fat diet (HFD) possess higher levels of bile acids, including conjugated bile acids, than mice fed a chow diet.^30^ Because increased substrate concentration may increase in vivo BSH activity, we also evaluated the ability of **AAA-10** to inhibit the enzyme activity in feces obtained from HFD-fed mice. We found that **AAA-10** inhibited BSH activity in this assay as effectively as **AAA-1**, our most potent first-generation BSH inhibitor (Figure S1B). Together, our data suggest that **AAA-10** is a potent inhibitor of BSHs found in the murine gut.

### AAA-10 inhibits recombinant BSHs and is more potent than AAA-2

To further characterize the potency of **AAA-10** compared to **AAA-1** and **AAA-2**, we determined the IC_50_ values of both **AAA-10** and **AAA-2** against purified recombinant *B. theta* (Gram-negative) and *B. longum* (Gram-positive) BSHs and compared these values to the IC_50_ values for **AAA-1** which had been determined in our previous work^24^ (Figure 3B-C). The IC_50_ values were evaluated using a conjugated bile acid substrate for which the enzymes demonstrated the best hydrolytic efficiency (TUDCA and TDCA, respectively).^24^ **AAA-10** exhibited an IC_50_ value of 10 nM against *B. theta* rBSH and 80 nM against *B. longum* rBSH, demonstrating that **AAA-10** was ∼250 fold more potent than **AAA-2** against *B. theta* rBSH and ∼15 fold more potent against *B. longum* rBSH (Table S1). Compared to **AAA-1, AAA-10** was ∼40 fold more potent against *B. theta* rBSH and equally potent against *B. longum* rBSH (Table S1). The increased potency of **AAA-10** against *B. theta* rBSH, a selective BSH, compared to both first-generation inhibitors highlights the potential of this compound to target BSHs which might otherwise be difficult to inhibit. These data demonstrate that we have developed a second-generation sulfonated inhibitor with increased potency compared to first-generation compounds.

### AAA-10 inhibits BSH activity in bacterial cultures

Next, we evaluated the ability of **AAA-10** to inhibit enzyme activity in growing cultures of BSH-containing bacteria using three Gram-negative and three Gram-positive strains found in the human gut (Figure 4A). Each bacterial culture was diluted to pre-log phase and co-incubated with 100 µM of **AAA-10** and 100 µM of an equimolar mixture of taurine-conjugated bile acids that are abundant in the murine gallbladder and small intestine (tauro-betamuricholic acid (TβMCA), taurocholic acid (TCA), TUDCA, and TDCA).^31^ Bacteria were then allowed to grow into stationary phase over 24 h. Because bacteria vary in their ability to metabolize different conjugated bile acids, this approach provides an unbiased way of testing the inhibitory activity of **AAA-10**. After 24 h, percent deconjugation was determined by quantifying bile acid concentrations in bacterial cultures by UPLC-MS. **AAA-10** exhibited near-complete inhibition of enzyme activity in all six bacterial cultures (<7% deconjugation) (Figure 4A, Figure S2). **AAA-10** displayed equivalent inhibitory activity to **AAA-1**, except in the case of *C. perfringens*, where **AAA-10** inhibited deconjugation to a greater extent than **AAA-1** (4% vs 22% deconjugation, respectively).^24^

**Figure 4.**
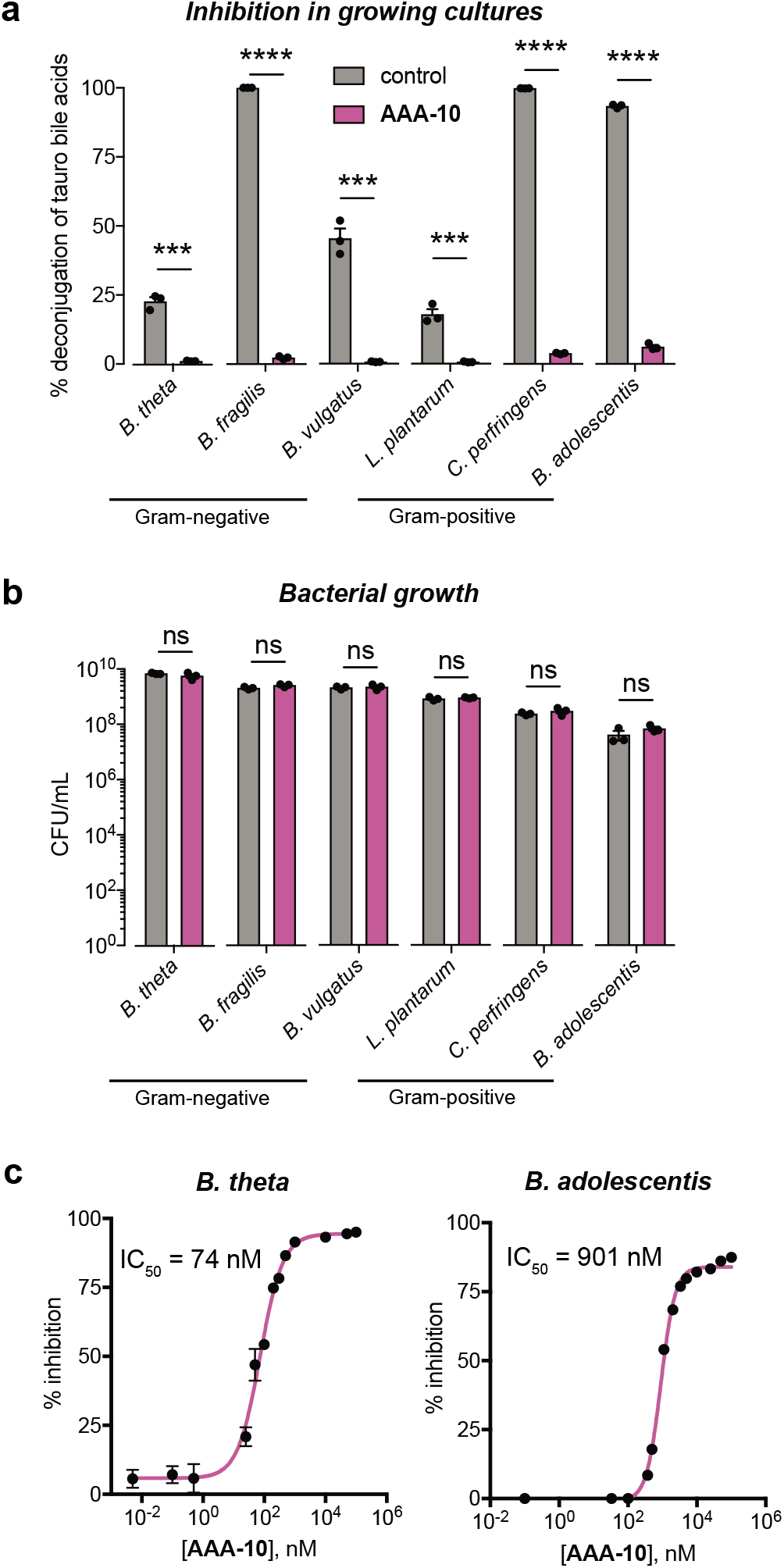
AAA-10 inhibits BSH activity in bacterial cultures without exhibiting antibacterial effects. **a, AAA-10** inhibits bacterial BSH activity. The BSH inhibitory activity of AAA-10 against three Gram-negative (*B. theta* VPI-5482, *Bacteroides fragilis* ATCC 25285, and *Bacteroides vulgatus* ATCC 8482) and three Gram-positive (*Lactobacillus plantarum* WCFS1, *Clostridium perfringens* ATCC 13124, and *Bifidobacterium adolescentis* L2–32) human gut bacteria was evaluated. BSH activity was quantified as percent deconjugation of tauro-conjugated bile acids at 24 h as determined by UPLC–MS (for absolute concentrations of substrates and products recovered, see Figure S2). **b, AAA-10** did not affect bacterial cell viability. At the end of the assay in (a), the bacteria were plated to determine cell viability. **c, AAA-10** is a nanomolar inhibitor of bacterial BSHs. Dose-response curves of **AAA-10** against *B. theta* and *B. adolescentis* cultures were generated using tauro-ursodeoxycholic acid (TUDCA) and tauro-deoxycholic acid (TDCA) as substrates, respectively. For **a**, and **b**, two-tailed Student’s t-test were performed. For **c**, Graphpad was used to fit IC_50_ curves. *p<0.05, **p<0.01, ***p<0.001, ****p<0.0001, ns = not significant. All assays were performed in biological triplicate, and data are presented as mean ± s.e.m.

To demonstrate that the BSH inhibitory activity of **AAA-10** in bacterial cultures was not due to growth inhibition, we evaluated the colony forming units in aforementioned bacterial cultures treated with **AAA-10**. We found that this compound did not significantly affect the growth of any of the tested bacterial strains at a concentration of 100 µM (Figure 4B).

We also determined the IC_50_ values of **AAA-10** against *B. theta* (Gram-negative) and *B. adolescentis* (Gram-positive) whole cell cultures. For this purpose, we used a single conjugated bile acid against which the enzyme demonstrated the highest deconjugation efficiency (TUDCA and TDCA, respectively). While the IC_50_ value for **AAA-10** against *B. adolescentis* was higher than the previously reported value for **AAA-1** (901 nM versus 108 nM, respectively), **AAA-10** displayed a lower IC_50_ value against *B. theta* than **AAA-1** (74 nM versus 427 nM, respectively)^24^ (Figure 4c). These data are consistent with our results using purified protein and show that **AAA-10** is the most potent inhibitor of the *B. theta* BSH yet developed. Together, these results demonstrate that **AAA-10** is a nanomolar inhibitor of gut bacterial BSHs that does not display anti-bacterial properties.

### AAA-10 displays limited off-target effects on mammalian cells

At high in vivo concentrations, bile acids have been shown to disrupt cell membranes and can induce apoptosis in mammalian cells^15,32,33^. Because **AAA-10** is based on a bile acid scaffold, we evaluated the toxicity of **AAA-10** on intestinal cells as well as its off-target effects on host bile acid receptors. Human intestinal Caco-2 cells were differentiated in transwell inserts to form a polarized monolayer with tight junctions^34^ (Figure S3A). Incubation of these cells with **AAA-1, AAA-2** or **AAA-10** showed that while **AAA-1** and **AAA-2** (100 µM) negatively affected the cell viability, **AAA-10** did not have an effect on the cell viability at this concentration (Figure 5A). **AAA-10** also had no effect on the viability of human liver cells (Hep-G2) at 100 µM or 500 µM concentrations (Figure S3B). We next determined whether BSH inhibitors affected intercellular tight junctions by measuring the passive diffusion of FITC-dextran (4 kDa) from the apical to the basolateral chamber of the transwells containing differentiated Caco-2 cells treated apically with our compounds. **AAA-10** did not appear to damage epithelial integrity at 100 µM or 500 µM concentrations, while **AAA-1, AAA-2**, and **AAA-9** increased FITC-d permeability by over 85% (∼1.5-3 fold) at a concentration of 100 µM (Figure 5B). In order to test the gut-restricted properties of **AAA-10**, we also quantified the amount of inhibitor in the apical and basolateral chambers in these transwell assays. We have previously shown that bile acids, including LCA, pass through Caco-2 monolayers.^35^ In contrast, while we were able to detect **AAA-10** in the apical chamber, no inhibitor was detected in the basolateral chamber 16 h after apical application, indicating that **AAA-10** does not pass through an epithelial monolayer (Figure 5C-D).

**Figure 5.**
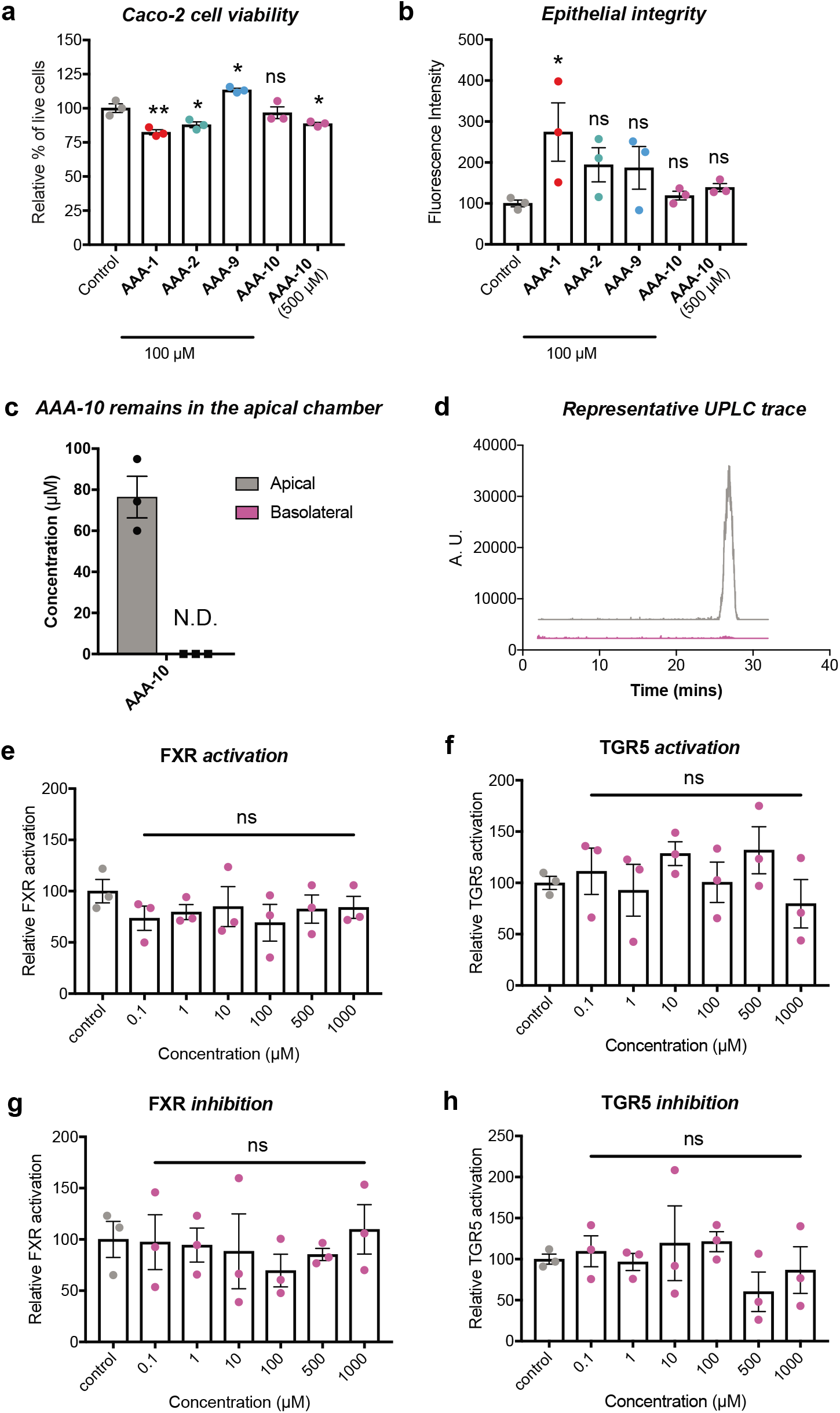
AAA-10 is not toxic to mammalian cells and not a ligand for FXR or TGR5. **a**, Incubation of differentiated Caco-2 cells with **AAA-10** (100 μM) did not result in toxicity, while incubation with an equivalent concentration of **AAA-1** and **AAA-2** resulted in decreased cell viability. **b, AAA-10** did not damage epithelial tight junctions at 100 μM or 500 μM, while treatment with **AAA-1** resulted in loss of epithelial integrity. Epithelial junction integrity was determined by measuring the transport of 4 kDa FITC-dextran from the apical to the basolateral chamber. **c, AAA-10** did not pass through an epithelial monolayer in an in vitro transwell assay (for assay setup see Figure S3A). Passage of the molecule from apical chamber to basolateral chamber was quantified by UPLC-MS. **d**, Representative UPLC-MS extracted ion chromatogram (EIC) traces of apical and basolateral chamber showing that no **AAA-10** was detected in the basolateral chamber. **e** and **f**, FXR and TGR5 agonist activity was measured by incubating Caco-2 cells with varying concentrations of **AAA-10** overnight. **g** and **h**, FXR and TGR5 antagonist activity was measured by incubating Caco-2 cells with varying concentrations of **AAA-10** overnight in the presence of 10 μM of the FXR agonist chenodeoxycholic acid (CDCA) or 10 μM of the TGR5 agonist lithocholic acid (LCA), respectively. For **a-b** and **e-h**, one-way ANOVA followed by Dunnett’s multiple comparisons test. *p<0.05, **p<0.01, ***p<0.001, ****p<0.0001, ns = not significant. All assays were performed in biological triplicate, and data are presented as mean ± s.e.m.

Bile acids can signal through the host receptors FXR and TGR5, thereby affecting host metabolism and immune function.^36^ Incubation of Caco-2 cells with increasing concentrations of **AAA-10** revealed that this compound did not act as an agonist of FXR or TGR5 (Figures 5E-F). Incubation of Caco-2 cells with either GW4062 (FXR agonist) or LCA (TGR5 agonist) followed by treatment with increasing concentrations of **AAA-10** revealed that this compound did not antagonize FXR or TGR5 (Figures 5G-H). Collectively, these data suggest that **AAA-10** is a potent pan-BSH inhibitor with low epithelial permeability that exhibits reduced off-target effects on host cells compared to the first-generation inhibitors **AAA-1** and **AAA-2**.

### AAA-10 reduces secondary bile acid abundance in vivo

We next evaluated the ability of **AAA-10** to inhibit BSH activity and modulate bile acid levels in vivo. Wild-type C57Bl/6J mice were gavaged once daily with **AAA-10** at a dose of 30 mg/kg for 5 days (Figure 6A). The inhibitor was administered at 6 pm to coincide with the start of the dark photoperiod when mice exhibit increased food consumption.^37^ Fecal BSH activity was significantly decreased on days 2 and 6 in **AAA-10**-treated mice compared to vehicle-treated mice (Figure 6B and Figure S4A), indicating that we were able to achieve durable BSH inhibition in vivo using **AAA-10**. Analysis of the cecal bile acid pool after sacrifice revealed that the abundances of DCA and LCA were significantly lowered in the **AAA-10**-treated group (Figures 6C). DCA and LCA are secondary bile acids that are produced exclusively by gut bacteria.^5^ Cecal **AAA-10** concentration was also negatively correlated with cecal concentrations of both DCA and LCA (Figures S4B-C). In addition, the abundances of DCA and LCA were decreased in feces each day starting on day 2 and overall in feces throughout the course of the study in **AAA-10**-treated mice compared to vehicle-treated mice (Figures 6D-E and Figure S4D). Together, these findings indicate that **AAA-10** treatment resulted in a sustained reduction in secondary bile acids in vivo over the period of study.

**Figure 6.**
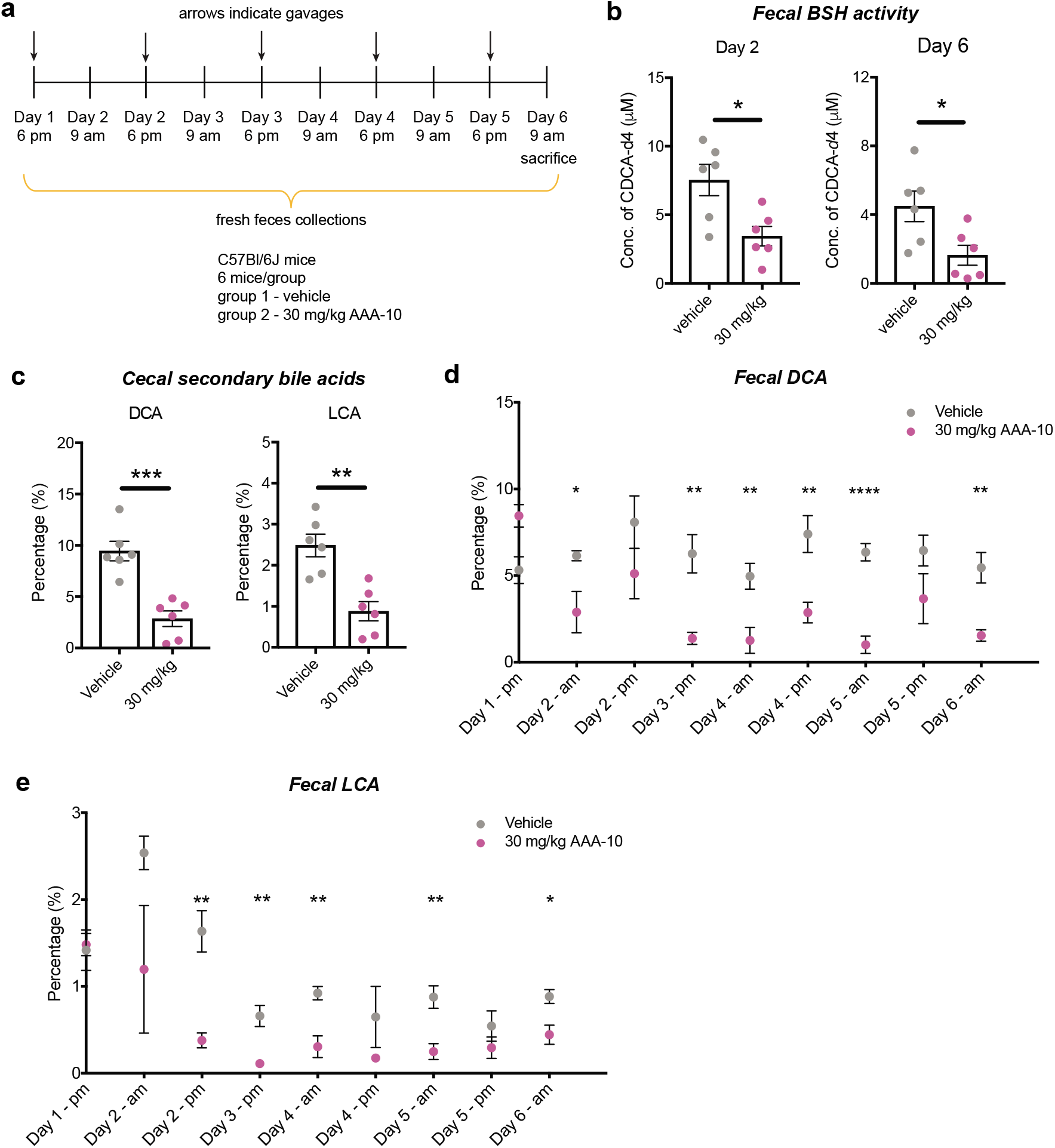
AAA-10 inhibits BSH activity in vivo and reduces secondary bile acid abundance. **a**, In vivo study design. C57Bl/6J mice were orally gavaged with **AAA-10 (**30 mg/kg) once daily for 5 days. Feces were collected daily and utilized to evaluate bile acid changes and BSH activity. Mice were sacrificed on day 6 and tissues and blood were collected. **b, AAA-10**-treated mice exhibited decreased BSH activity compared to vehicle-treated mice in fresh feces collected on days 2 and 6. **c**, Percentages of the secondary bile acids deoxycholic acid (DCA) and lithocholic acid (LCA) were reduced in cecal contents of mice treated with **AAA-10. d** and **e**, Analysis of fecal bile acid contents over the period of the study showed that abundances of the two secondary bile acids DCA and LCA were consistently decreased throughout the experiment. For **b-e**, n=6 mice/group, two-tailed Welch’s t test was performed. *p<0.05, **p<0.01, ***p<0.001, ****p<0.0001, ns = not significant. All data are presented as mean ± s.e.m.

Finally, to evaluate the gut permeability of **AAA-10**, we quantified the levels of this compound in cecal contents and plasma at sacrifice and in feces over the course of the experiment (Figures S4E-G). We observed a mean value of 276 picomol/mg wet mass (∼276 μM) of **AAA-10** in cecal contents and a range of 66-2087 picomol/mg (∼66-2087 μM) in feces. In contrast, five of the six mice exhibited undetectable levels of **AAA-10** in plasma (Figure S4G). In a separate experiment in which mice were sacrificed 4 hours after the final gavage, we detected mean **AAA-10** concentrations of 128 picomol/mg wet mass (∼128 μM) in cecal contents and 12 nM in plasma (Figures S4H-I). Together, these data indicate that AAA-10 displays high colonic exposure and low gut permeability.

## CONCLUSION

The studies reported herein were initiated with the goal of improving the potency of the first-generation gut-restricted inhibitor **AAA-2**. Structure-activity relationship studies that focused on incorporating the carbon scaffolds of different abundant bile acids into our inhibitor design led to the identification of a second-generation inhibitor, **AAA-10**. This compound exhibited increased potency in an array of in vitro assays compared to **AAA-2**. The structure of **AAA-10** is based on the core of LCA, a bile acid that contains a single hydroxyl group at C3. In previous work, we characterized the BSH activity of 20 abundant human gut Bacteroidetes species against glyco- and tauro-conjugated bile acids.^20^ Glyco-lithocholic acid (GLCA) and tauro-lithocholic acid (TLCA) were effectively deconjugated (>90% conversion) by all BSH-containing Bacteroidetes species tested. In contrast, all other bile acids were incompletely deconjugated (<70%) by two or more of the species tested. Taken together, these results suggest that the LCA core may be effective as a scaffold for BSH inhibitors because it is recognized as a substrate by a range of gut bacteria. Future studies testing the deconjugating ability of a variety of Gram-positive and Gram-negative strains against a panel of conjugated bile acid substrates may further elucidate the substrate scope of gut bacterial BSHs and thus aid in next-generation inhibitor design.

**AAA-10** also exhibited an improved off-target effects profile compared to **AAA-1** and **AAA-2**. This compound inhibited BSH activity without inhibiting the growth of the Gram-negative and Gram-positive bacteria tested, did not activate or inhibit the host bile acid receptors FXR or TGR5 at concentrations expected to be effective in vivo, and was found to be non-toxic to intestinal and liver cells at higher concentrations than **AAA-1** and **AAA-2**.

Finally, we demonstrated that **AAA-10** inhibits BSH activity in vivo. Once-daily administration of **AAA-10** by gavage at a dose of 30 mg/kg resulted in significant BSH inhibition in feces and a decrease in the abundance of the secondary bile acids DCA and LCA in feces and cecal contents. DCA and LCA are known to play crucial roles in host physiology. On the one hand, DCA and LCA are strongly associated with colon cancer development in patients, and evidence indicates that these compounds promote carcinogenesis in the colon and liver.^13,14,38-40^ On the other hand, DCA has been shown to limit growth of the pathogen *Clostridium difficile*,^41^ and recent work has shown that LCA induces the production of the anti-diabetic metabolite cholic acid-7-sulfate.^35^ The ability to modulate the abundance of these compounds in vivo in fully colonized animals will facilitate investigations of the roles of these molecules in host physiology.

Looking ahead, it will be valuable to consider whether long-term use of BSH inhibitors in vivo affects microbial community composition. In addition, once-daily dosing via oral gavage may not be optimal in the case of ad libitum feeding, and further optimization of a strategy for inhibitor administration may be required. Nonetheless, our data indicate that we have developed a potent, non-toxic BSH inhibitor that modulates the in vivo bile acid pool, shifting the bile acid pool away from DCA and LCA. Importantly, we have shown that **AAA-10** exhibits in vivo efficacy using a daily dosing strategy in a 5 day experiment, a finding that paves the way for the use of **AAA-10** in longer-term animal models to elucidate how bile acids affect host health and either drive or ameliorate disease phenotypes. Because bile acids are absorbed and recirculated to the liver via the portal vein,^5,42^ BSH inhibitors will facilitate investigations of how bile acids are causally involved in the initiation and progression of both liver and GI tract disorders, including inflammatory bowel diseases, non-alcoholic fatty liver disease (NAFLD), non-alcoholic steatohepatitis (NASH), liver cirrhosis, and liver and colon cancer.^17,19^ Demonstration of prevention or amelioration of disease phenotypes in animals would suggest that BSHs could be targeted in a therapeutic context to treat human disease.

## METHODS

Full details for all materials and methods are provided in the Supporting Information.

## ASSOCIATED CONTENT

### Supporting Information

Materials and methods (including detailed synthetic protocols and characterization data), Figures S1-4, Table S1, Scheme S1.

## AUTHOR INFORMATION

### Author Contributions

A.A.A. and A.S.D. conceived the project and designed the experiments. A.A.A. performed the synthesis and most of the experiments. D. R. and A.S.B. performed the in vivo BSH inhibition and bile acid pool modulation study. S.N.C. performed the cell culture assays. C.E.P. performed the experiments with Gram-negative bacteria. M.D.M. heterologously expressed and purified *B. longum* recombinant BSH. A.A.A. and A.S.D. wrote the manuscript. All authors edited and contributed to the review of the manuscript.

### Funding

This research was supported National Institutes of Health (NIH) grant R35 GM128618 (A.S.D), an Innovation Award from the Center for Microbiome Informatics and Therapeutics at MIT (A.S.D), a grant from Harvard Digestive Diseases Center (supported by NIH grant 5P30DK034854-32) (A.S.D), a John and Virginia Kaneb Fellowship (A.S.D), a Quadrangle Fund for the Advancement and Seeding of Translational Research at Harvard Medical School (Q-FASTR) grant (A.S.D), and an HMS Dean’s Innovation Grant in the Basic and Social Sciences (A.S.D). S.N.C. acknowledges an American Heart Association Postdoctoral Fellowship. M.D.M. acknowledges an NSF Graduate Research Fellowship (DGE1745303).

### Notes

The authors declare the following competing financial interest(s): A. Sloan Devlin is an ad hoc consultant for Takeda Pharmaceuticals and HP Hood. The other authors declare that no competing interests exist.

## ACKNOWLEDGEMENTS

We are indebted to Stephen Blacklow, Nathanael Gray, David Scott, John M. Hatcher, Jinhua Wang, Jon Clardy, and members of the Devlin and Clardy groups for helpful discussions. We would like to acknowledge the Blacklow and Kruse labs for help with equipment and reagents. We thank K. Schoonjans (Ecole polytechnique fédérale de Lausanne-EPFL) for the FXR reporter plasmid and the ICCB-Longwood Screening Facility for use of their fluorescent plate reader. We thank the scientists at Bienta, Enamine for help with in vivo pharmacokinetics experimental design and execution.

## REFERENCES

1. Donia, M. S. & Fischbach, M. A. Small molecules from the human microbiota. Science 349, 1254766 (2015).

2. Wallace, B. D. et al.. Alleviating cancer drug toxicity by inhibiting a bacterial enzyme. Science 330, 831–835 (2010).

3. Roberts, A. B. et al.. Development of a gut microbe-targeted nonlethal therapeutic to inhibit thrombosis potential. Nat. Med. 24, 1407–1417 (2018).

4. Maini Rekdal, V., Bess, E. N., Bisanz, J. E., Turnbaugh, P. J. & Balskus, E. P. Discovery and inhibition of an interspecies gut bacterial pathway for Levodopa metabolism. Science 364, eaau6323 (2019).

5. Ridlon, J. M., Kang, D.-J. & Hylemon, P. B. Bile salt biotransformations by human intestinal bacteria. J. Lipid Res. 47, 241–259 (2006).

6. Hofmann, A. F. The function of bile salts in fat absorption. The solvent properties of dilute micellar solutions of conjugated bile acids. Biochem. J. 89, 57–68 (1963).

7. Roda, A., Hofmann, A. F. & Mysels, K. J. The influence of bile salt structure on self-association in aqueous solutions. J. Biol. Chem. 258, 6362–6370 (1983).

8. Fiorucci, S. & Distrutti, E. Bile Acid-Activated Receptors, Intestinal Microbiota, and the Treatment of Metabolic Disorders. Trends Mol Med 21, 702–714 (2015).

9. Schaap, F. G., Trauner, M. & Jansen, P. L. M. Bile acid receptors as targets for drug development. Nat Rev Gastroenterol Hepatol 11, 55–67 (2014).

10. Pols, T. W. H. et al.. Lithocholic acid controls adaptive immune responses by inhibition of Th1 activation through the Vitamin D receptor. PLOS ONE 12, e0176715 (2017).

11. Hang, S. et al.. Bile acid metabolites control TH17 and Treg cell differentiation. Nature 576, 143–148 (2019).

12. Begley, M., Hill, C. & Gahan, C. G. M. Bile salt hydrolase activity in probiotics. Appl. Environ. Microbiol. 72, 1729–1738 (2006).

13. Ridlon, J. M., Wolf, P. G. & Gaskins, H. R. Taurocholic acid metabolism by gut microbes and colon cancer. Gut Microbes 7, 201–215 (2016).

14. Ma, C. et al.. Gut microbiome-mediated bile acid metabolism regulates liver cancer via NKT cells. Science 360, eaan5931 (2018).

15. Ajouz, H., Mukherji, D. & Shamseddine, A. Secondary bile acids: an underrecognized cause of colon cancer. World J Surg Oncol 12, 164–5 (2014).

16. Ridlon, J. M., Harris, S. C., Bhowmik, S., Kang, D.-J. & Hylemon, P. B. Consequences of bile salt biotransformations by intestinal bacteria. Gut Microbes 7, 22–39 (2016).

17. Song, Z. et al.. Taxonomic profiling and populational patterns of bacterial bile salt hydrolase (BSH) genes based on worldwide human gut microbiome. Microbiome 7, 9 (2019).

18. Parasar, B. et al.. Chemoproteomic Profiling of Gut Microbiota-Associated Bile Salt Hydrolase Activity. ACS Cent Sci 5, 867–873 (2019).

19. Jia, B., Park, D., Hahn, Y. & Jeon, C. O. Metagenomic analysis of the human microbiome reveals the association between the abundance of gut bile salt hydrolases and host health. Gut Microbes 11, 1300–1313 (2020).

20. Yao, L. et al.. A selective gut bacterial bile salt hydrolase alters host metabolism. eLife 7, 675 (2018).

21. Joyce, S. A. et al.. Regulation of host weight gain and lipid metabolism by bacterial bile acid modification in the gut. Proc. Natl. Acad. Sci. U.S.A. 111, 7421–7426 (2014).

22. Li, F. et al.. Microbiome remodelling leads to inhibition of intestinal farnesoid X receptor signalling and decreased obesity. Nat Commun 4, 2384 (2013).

23. Xie, C. et al.. An Intestinal Farnesoid X Receptor-Ceramide Signaling Axis Modulates Hepatic Gluconeogenesis in Mice. Diabetes 66, 613–626 (2017).

24. Adhikari, A. A. et al.. Development of a covalent inhibitor of gut bacterial bile salt hydrolases. Nature Chemical Biology 16, 318–326 (2020).

25. Tanaka, H., Hashiba, H., Kok, J. & Mierau, I. Bile salt hydrolase of Bifidobacterium longum-biochemical and genetic characterization. Appl. Environ. Microbiol. 66, 2502– 2512 (2000).

26. Wang, Z. et al.. Identification and Characterization of a Bile Salt Hydrolase from Lactobacillus salivarius for Development of Novel Alternatives to Antibiotic Growth Promoters. Appl. Environ. Microbiol. 78, 8795–8802 (2012).

27. Strott, C. A. Sulfonation and molecular action. Endocr Rev 23, 703–732 (2002).

28. Alnouti, Y. Bile Acid sulfation: a pathway of bile acid elimination and detoxification. Toxicol. Sci. 108, 225–246 (2009).

29. Palmer, J. T. & Inc, P. Process for forming a fluoromethyl ketone. (1994).

30. Zheng, X. et al.. Bile acid is a significant host factor shaping the gut microbiome of diet-induced obese mice. BMC Biol 15, 120–15 (2017).

31. Sayin, S. I. et al.. Gut microbiota regulates bile acid metabolism by reducing the levels of tauro-beta-muricholic acid, a naturally occurring FXR antagonist. Cell Metab. 17, 225– 235 (2013).

32. Perez, M.-J. & Briz, O. Bile-acid-induced cell injury and protection. World J. Gastroenterol. 15, 1677–1689 (2009).

33. Glinghammar, B., Inoue, H. & Rafter, J. J. Deoxycholic acid causes DNA damage in colonic cells with subsequent induction of caspases, COX-2 promoter activity and the transcription factors NF-kB and AP-1. Carcinogenesis 23, 839–845 (2002).

34. Ferruzza, S., Rossi, C., Scarino, M. L. & Sambuy, Y. A protocol for differentiation of human intestinal Caco-2 cells in asymmetric serum-containing medium. Toxicol In Vitro 26, 1252–1255 (2012).

35. Chaudhari, S. N. et al.. A microbial metabolite remodels the gut-liver axis following bariatric surgery. Cell Host & Microbe 317, 571 (2020).

36. Wahlström, A., Sayin, S. I., Marschall, H.-U. & Bäckhed, F. Intestinal Crosstalk between Bile Acids and Microbiota and Its Impact on Host Metabolism. Cell Metab. 24, 41–50 (2016).

37. Ellacott, K. L. J., Morton, G. J., Woods, S. C., Tso, P. & Schwartz, M. W. Assessment of feeding behavior in laboratory mice. Cell Metab. 12, 10–17 (2010).

38. Reddy, B. S., Narasawa, T., Weisburger, J. H. & Wynder, E. L. Promoting effect of sodium deoxycholate on colon adenocarcinomas in germfree rats. J. Natl. Cancer Inst. 56, 441–442 (1976).

39. Narisawa, T., Magadia, N. E., Weisburger, J. H. & Wynder, E. L. Promoting effect of bile acids on colon carcinogenesis after intrarectal instillation of N-methyl-N’-nitro-N-nitrosoguanidine in rats. J. Natl. Cancer Inst. 53, 1093–1097 (1974).

40. Yoshimoto, S. et al.. Obesity-induced gut microbial metabolite promotes liver cancer through senescence secretome. Nature 499, 97–101 (2013).

41. Buffie, C. G. et al.. Precision microbiome reconstitution restores bile acid mediated resistance to Clostridium difficile. Nature 517, 205–208 (2015).

42. van de Peppel, I. P., Verkade, H. J. & Jonker, J. W. Metabolic consequences of ileal interruption of the enterohepatic circulation of bile acids. Am. J. Physiol. Gastrointest. Liver Physiol. 319, G619–G625 (2020).

