## Supporting Information for "A gut-restricted lithocholic acid analog as an inhibitor of gut bacterial bile salt hydrolases"

Materials and Methods ........................................................ 2-22

### Materials and Methods

**Reagents.** All bile acids and reagents for synthesis were commercially purchased from Steraloids Inc. and Sigma Aldrich. Stock solutions of all bile acids and inhibitors were prepared in molecular biology grade DMSO (Sigma Aldrich) at 1000X concentrations. Solvents used for preparing UPLC-MS samples were HPLC grade. New biological materials reported here are available from the authors upon request. 100 μM bile acid pool consisted of TCA, TβMCA, TUDCA and TDCA (25 μM each).

**Bacterial culturing.** All bacterial strains were cultured at 37 °C in $BHI^+$ (Bacto™ BHI, BD, supplemented with 5 mg $l^{-1}$ hemin, and 2.5 ul $l^{-1}$ Vitamin $K_1$). All strains were grown under anaerobic conditions in an anaerobic chamber (Coy Lab Products Airlock) with a gas mix of 5% hydrogen and 20% carbon dioxide nitrogen. *Escherichia coli* was grown aerobically at 37 °C in LB medium supplemented with ampicillin to select for the pET21b plasmid.

**UPLC-MS analysis.** Bile acid profiling by UPLC-MS was performed using a published method.[1] Correction factors for extraction efficiency were used and were determined by extraction of known concentrations of relevant bile acids from buffer or bacterial media and comparison to standard curves. The limits of detection for individual bile acids were determined using commercially available standards/ synthesized compounds solubilized in 1:1 MeOH/water and are as follows: βMCA, 0.03 picomol $\mu l^{-1}$; TβMCA, 0.01 picomol $\mu l^{-1}$; CA, 0.04 picomol $\mu l^{-1}$; TCA, 0.01 picomol $\mu l^{-1}$; UDCA, 0.04 picomol $\mu l^{-1}$; TUDCA, 0.01 picomol $\mu l^{-1}$; DCA, 0.04 picomol $\mu l^{-1}$; TDCA, 0.05 picomol $\mu l^{-1}$; GCDCA-d4, 0.1 picomol $\mu l^{-1}$; CDCA-d4, 0.1 picomol $\mu l^{-1}$; 7-oxo-CA, 0.5 picomol $\mu l^{-1}$; **AAA-2**, 0.05 picomol $\mu L^{-1}$; **AAA-10**, 1.0 picomol $\mu l^{-1}$.

**Screen of inhibitors in conventional mouse feces.** BSH activity in fecal pellets was quantified using a modified version of a published method.[2,3] Fresh feces were collected from wildtype adult male C57Bl6/J mice. Fecal pellets (approximately 10-20 mg) were broken into fine particles in buffer (PBS with 0.25 mM TCEP) to obtain a concentration of 1 mg $ml^{-1}$. Indicated concentrations of inhibitors were added to the fecal slurry and the mixture was incubated at 37 °C for 30 mins. 100 μM glycochenodeoxycholic acid-d4 (GCDCA-d4) was added to the mixture and incubated at 37 °C for 18 h. The tubes were then frozen in dry ice for 5 mins and upon thawing were diluted with an equal volume of HPLC grade methanol. The slurry was centrifuged at 12,500 g for 10 mins. The supernatant was removed into a clean Eppendorf tube and centrifuged again. The supernatant was transferred to MS vials and samples were analyzed as per the method described in "UPLC-MS Analysis". The concentration of product detected from these assays was reported directly.

**Protein expression and purification.** *B. thetaiotaomicron* and *B. longum* recombinant BSHs were expressed and purified as previously described.[3]

**Determination of $IC_{50}$ values of inhibitors against recombinant proteins.** 200 nM rBSH was incubated with increasing concentrations of the inhibitors at 37 °C for 1 h in 1 ml PBS buffer containing 0.25 mM TCEP and 5% glycerol at pH 7.5. 100 μM bile acid (TUDCA for *B. theta* BSH and TDCA for *B. longum* BSH) was added to the above solution and incubated at 37 °C for 2 h. The solution was acidified to pH = 1 using 6M HCl and extracted twice with 1 ml ethyl acetate. The combined organic layers were then dried using a Biotage TurboVap LV. The dried extracts were resuspended in 1:1 methanol:water and transferred to mass spectrometry vials.

Samples were analyzed as per the method described in UPLC–MS analysis. The obtained concentrations of bile acids were used to determine percentage deconjugation.

**Equation for calculating percent deconjugation.**

Percent deconjugation = concentration of deconjugated bile acids detected / (concentration of deconjugated bile acids detected + concentration of conjugated bile acids detected) X 100.

**Equation for calculating percent inhibition.**

Percent inhibition = (percent deconjugation in control sample - percent deconjugation in inhibitor-treated sample) / percent deconjugation in control sample X 100.

**BSH inhibition in bacterial cultures.** Bacterial cultures were diluted to $OD_{600}$ of 0.1 in 4 ml $BHI^+$, containing 100 μM taurine conjugated bile acid pool and 100 μM **AAA-10**. These cultures were then grown anaerobically at 37 °C. After 21 h, serial dilutions were plated on $BHI^+$ agar to determine cell viability (CFU $ml^{-1}$). 1 ml of the entire bacterial culture was acidified to pH = 1 using 6M HCl followed by addition of 2 ml ethyl acetate and vortexed. The cultures were spun down in a centrifuge at 2,500 g for 5 mins to obtain better separation. The organic layer was then removed and the aqueous layer was extracted again using 2 ml of ethyl acetate. The dried organic extracts were resuspended in 1:1 methanol:water and transferred to mass spec vials and analyzed as per the method described in "UPLC-MS Analysis". The obtained concentrations of bile acids were used to determine percent deconjugation.

**Determination of $IC_{50}$ values of AAA-10 in bacterial cultures.** Note that due to slow growth of *B. longum*, *B. adolescentis* was used as a representative of Gram-positive bacteria. Overnight cultures of *B. theta* and *B. adolescentis* were diluted to an $OD_{600}$ of 0.1 in 2 ml fresh CHG media (see "Bacterial Culturing") containing 100 μM TUDCA or TDCA, respectively, and inhibitor at increasing concentrations. *B. theta* and *B. adolescentis* deconjugated TUDCA and TDCA, respectively, to the greatest extent of any of the conjugated substrates, and therefore these substrates were used to determine $IC_{50}$ values. Cultures were then grown anaerobically at 37 °C for 24 h (*B. adolescentis*) or 48 h (*B. theta*). Longer incubation time was required for *B. theta* because for this bacterium, significant BSH activity was only observed during stationary phase. Cultures were extracted and analyzed as per the method described in "BSH Inhibition in Bacterial Cultures" to determine percent deconjugation which was then converted into percent inhibition.

**Cell culture.** Caco-2 cells and HepG2 cells were obtained from American Type Culture Collection (Manassas, VA). Caco-2 cells were maintained in Minimum Essential Medium (MEM) supplemented with GlutaMAX. All cell culture media was supplemented with 10% fetal bovine serum (FBS), 100 units $ml^{-1}$ penicillin, and 100 μg $ml^{-1}$ streptomycin (GenClone). Cells were grown in FBS- and antibiotic-supplemented 'complete' media at 37 °C in an atmosphere of 5% $CO_2$.

**Plasmids and transient transfections.** For luciferase reporter assays, vectors expressing human reporter constructs were used. The pGL4.29[luc2P/CRE/Hygro] plasmid (Promega Corporation) and the pGL4 [Shp-luc] plasmid obtained from Kristina Schoonjans lab at EPLF was transiently transfected in Caco-2 cells at a concentration of 2 μg $ml^{-1}$ of media each for studying TGR5 and FXR activation respectively. The pGL4.74[hRluc/CMV] plasmid (Promega Corporation) was used as a transfection efficiency control at a concentration of 0.05 μg $ml^{-1}$ of media. All plasmids were transfected using Opti-MEM (Gibco) and Lipofectamine 2000 (Invitrogen, Life Technologies, Grand Island, NY, USA) according to manufacturer's instructions. After overnight incubation, **AAA-10** and/or bile acids were added in complete media. **AAA-10** and/or bile acids were diluted in DMSO and the concentration of DMSO was kept constant. 10 μM of LCA or 10 μM of CDCA was added along with **AAA-10** to study TGR5 and FXR antagonism respectively and incubated overnight. Cells were harvested the next day for the luciferase assay.

**Luciferase reporter assay.** Luminescence was measured using the Dual-Luciferase Reporter Assay System (Promega Corporation) according to manufacturer's instructions. Cells were washed gently with PBS and lysed in PLB from the kit. Luminescence was measured using a SpectraMax M5 plate reader (Molecular Devices, San Jose, CA) at the ICCB-Longwood Screening Facility at HMS. Luminescence was normalized to *Renilla* luciferase activity and percentage relative luminescence was calculated compared to DMSO control.

**Epithelial permeability assay.** Undifferentiated Caco-2 cells were seeded in 24-well plate transwells (0.4 μM pore size, Costar) at 200,000 cells per transwell. Media was changed on days 4, 8, 12, 16, and 18 to differentiate Caco-2 cells in vitro.[4] On day 21, fully differentiated and polarized cells were used for FITC-dextran permeability assay. Briefly, inhibitors were added in PBS at indicated concentrations to the apical chamber of the transwells containing differentiated Caco-2 cells and incubated overnight. The apical chamber of the transwells contained a volume of 100 μl PBS with inhibitor or DMSO control, while the basolateral chamber contained 500 $\mu$l of PBS. Caco-2 epithelial integrity was assayed by measuring passive diffusion of 4 kDa FITC-Dextran (Sigma Aldrich) added at a concentration of 5 μM to the apical chamber. Diffusion from the apical to basolateral side was measured by fluorescence reading in PBS on the basolateral side of the transwell system using a SpectraMax M5 plate reader (Molecular Devices, San Jose, CA) at the ICCB-Longwood Screening Facility at HMS. Fluorescence reading was normalized to the DMSO control. Transport of inhibitors from the apical to basolateral compartment was measured by drying basolateral media under vacuum and resuspending contents in methanol prior to injecting in the UPLC-MS.

**Cell viability assay.** Cells were treated with indicated compounds diluted in DMSO in complete MEM media. The viability of differentiated Caco-2 cells in transwells was measured using an MTT assay (Abcam, ab211091), while the viability of HepG2 cells was measured using Cell Titer Glo (Promega) according to manufacturer's instructions. The concentration of DMSO was kept constant and used as a negative control. Cells were incubated with the compounds overnight at 37 °C in an atmosphere of 5% CO2. The next day, cell viability was measured. For differentiated Caco-2 cells, cell culture media was replaced with the MTT reagent and incubated for 3 hours at 37 °C. Following incubation, the MTT reagent was replaced with the MTT solvent and incubated for 15 min, followed by colorimetric analysis. The SpectraMax M5 plate reader (Molecular Devices, San Jose, CA) at the ICCB-Longwood Screening Facility at HMS was used to measure cell viability via luminescence (Cell Titer Glo, HepG2 cells) or absorbance (MTT reagent, Caco-2 cells) at 690 nm. Percentage relative viability was calculated compared to DMSO control.

**BSH inhibition and bile acid pool modulation in vivo.** Male C57BL/6J mice obtained from Jackson laboratories were maintained under a strict 12 h/12 h light/dark cycle and under constant temperature (21 ± 1 °C) and humidity (55–65%). All experiments were conducted on 13-14 week-old mice. Mice were maintained on a standard chow diet (Purina LabDiet, catalog no. 5008) for the duration of the experiment. Mice were split into two groups of six mice each and were gavaged once daily at the beginning of the dark phase with either 200 μl of 95% PBS/ 5% DMSO containing 10% captisol (w/v) (vehicle group) or with 200 μl of 95% PBS/ 5% DMSO containing 10% captisol (w/v) and AAA-10 at a concentration of 3.75 mg ml-1 (treatment group) for five consecutive days. For the fecal pellet collection, each mouse was transferred to a temporary cardboard cage for a several minutes until a fecal pellet was produced.

**BSH activity in feces.** BSH activity in fecal pellets were quantified using a modified version of a published method.[2,3] Fecal pellets (approximately 10-20 mg) were suspended in buffer (PBS with 0.25 mM TCEP) containing 100 μM (GCDCA-d4) to obtain a concentration of 20 mg $ml^{-1}$. The fecal pellets were broken into fine particles and the mixture was incubated at 37 °C for 25 mins. Samples were processed and analyzed as per the method described in "Screen of Inhibitors in Conventional Mouse Feces". The concentration of product detected from these assays were reported directly.

**Quantification of bile acids and AAA-10 in tissues and plasma.** Bile acids and AAA-10 were extracted from tissues and plasma and quantified using a previously published method.[1]

**Determination of bile acid abundance in feces over the entire period of the study.** The following formula was used to determine the abundance of a particular bile acid over the entire period of the study:

Bile acid abundance over the study = (Picomol of a bile acid of interest at timepoint 1 + picomol of a bile acid of interest at timepoint 2 + …. + picomol of a bile acid of interest at last timepoint) / (total picomol of all fecal bile acids at timepoint 1 + total picomol of all fecal bile acids at timepoint 2 + …. + total picomol of all fecal bile acids at last timepoint). The obtained value was then divided by the total amount of bile acids detected over the period of study * 100.

**In vivo pharmacokinetics of AAA-10 at 4h timepoint.** C57BL/6 mice obtained from the facility at Bienta Enamine were maintained under a strict 12 h/12 h light/dark cycle and a constant temperature (22 ± 3 °C) and humidity (40–70%). All experiments were conducted on

10-11 week old male mice. During a seven day of acclimatization period, each animal was kept in a separate cage and had limited access to food (free access twice a day, from 9:00 h to 13:00 h and from 18:00 h to 22:00 h). Mice had no access to food at other times. Mice had free access to acidified boiled tap water. All animals were under observation and only animals without any clinical symptoms of illness were taken into the study. During treatment with **AAA-10,** mice were fed according to the above schedule and treatment was carried out 2 times a day immediately before providing access to food (at 9:00 h and 18:00 h) on the days 1-4 of the experimental period and once a day (9:00 h, before providing access to food) on the terminal ($5^{th}$) day of the experimental period. Treatment group animals were gavaged with 150 μl of PBS containing 10% captisol (w/v) and **AAA-10** at a concentration of 3.75 mg $ml^{-1}$ while the control group animals were treated with 150 μl of PBS containing 10% captisol (w/v). Plasma and tissue sample collections (liver, kidney, heart, duodenum, jejunum, ileum, cecum and colon) were performed at the terminal sacrifice on the $5^{th}$ day of the study four hours after the final gavage.

**Ethics.** For the BSH inhibition in vivo studies, mouse experiments were performed under the approval of the Beth Israel Deaconess Medical Center IACUC. For the in vivo pharmacokinetics studies, protocols were approved by Bienta's Animal Care and Use Committee and experiments were performed in adherence with the European Convention for the Protection of Vertebrate Animals used for Experimental and other Scientific Purposes (1986). BACUC protocol number - №9-3/2020 dated 30.09.2020.

**Code availability statement**

No custom code or mathematical algorithms were used in this study.

### Synthetic procedures

**General:** All anhydrous reactions were run under a positive pressure of argon or nitrogen. Anhydrous methylene chloride (DCM) and tetrahydrofuran (THF) were purchased from Sigma Aldrich. Silica gel column chromatography was performed using 60 Å silica gel (230−400 mesh). NMR spectra recorded in $CDCl_3$ used residual chloroform or TMS as the internal reference.

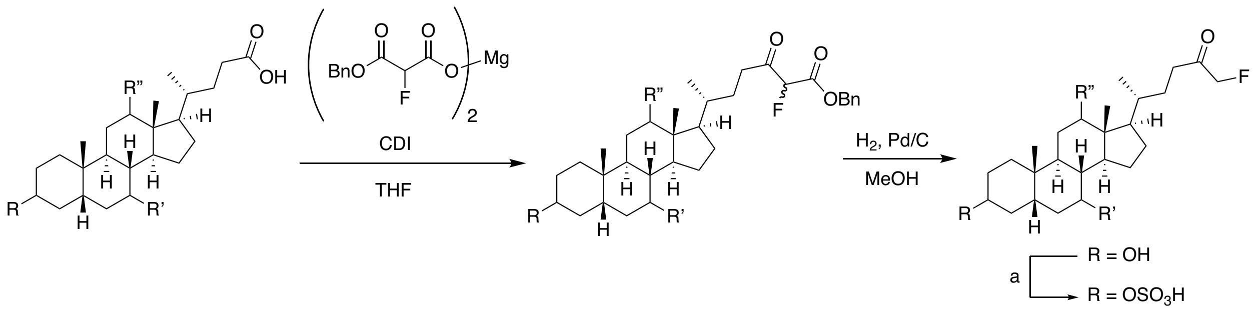

**Scheme 1: Generic scheme for synthesis of compounds 1-5.** CDI = carbonyldiimidazole, a = $SO_3$.pyridine, pyridine

**General procedure for the synthesis of α-fluoromethyl ketone analogs.**

The magnesium benzyl fluoromalonate coupling reagent was synthesized according to a reported protocol[5] with modifications.

Step 1. To a 0.3 M solution of the C-24 acid (1.0 equiv.) in anhydrous THF, 1'-carbonyldiimidazole (CDI) (1.0 equiv.) was added and stirred at rt for 1 h. The magnesium benzyl fluoromalonate (2.0 equiv.) was suspended in anhydrous THF (0.3 M) and the above solution was added dropwise. The resultant mixture was stirred at rt for 18 h. The reaction was quenched by the addition of 10 mL of 1M HCl and concentrated using a rotary evaporator. The residue was then partitioned using 10 mL ethyl acetate and 10 mL water. The organic layer was separated and the aqueous layer was extracted with ethyl acetate (2 x 10 mL). The combined organic layers were then dried over sodium sulfate, filtered and concentrated. The crude compound was then purified by silica gel chromatography (20% ethyl acetate/80% hexanes).

<u>Step 2.</u> To a 0.15 M solution of above compound (1.0 equiv.) in methanol, palladium on carbon (0.05 equiv.) was added. The flask was vacuumed and replaced with a hydrogen balloon. The reaction mixture was stirred at rt for 3 h. The solution was then filtered through a celite bed and the filtrate was concentrated to provide the crude compound. The compound was then purified by silica gel chromatography using a gradient of 50 to 60 % ethyl acetate in hexanes to provide the pure compound.

**AAA-1 (**compound 7; **1**). Synthesis and characterization previously described. [3]

**AAA-3** (**9**)**.** Yield = 97 mg (19% over 2 steps); TLC $R_f$ = 0.33 (30% ethyl acetate/70% hexanes); $^1$H NMR (400 MHz, $CDCl_3$) δ 4.79 (d, *J* = 47.6 Hz, 2H), 3.97 (br s, 1H), 3.565-3.57 (m, 1H), 2.63-2.43 (m, 2H), 1.91-1.23 (m, 23H), 1.18-0.94 (m, 6H), 0.91 (s, 3H), 0.68 (s, 3H); $^{13}$C NMR (100 MHz, $CDCl_3$) δ 207.53 (d, *J* = 19.2 Hz), 84.91 (d, *J* = 184.3), 73.11 (d, *J* = 11.2 Hz), 71.75, 48.28, 47.21, 46.48, 42.06, 36.44, 36.03, 35.18, 35.12, 34.95, 34.10, 33.69, 30.51, 28.75, 28.59, 27.40, 26.10, 23.60, 23.14, 17.43, 12.74, 12.73; HRMS (*m/z*): [M + HCOO - H]$^-$ calcd. for $C_{26}H_{42}FO_5$, 453.3022; found, 453.3030.

**AAA-5** (**10**)**.** Yield = 142 mg (27% over 2 steps); TLC $R_f$ = 0.44 (30% ethyl acetate/70% hexanes); $^1$H NMR (400 MHz, $CDCl_3$) δ 4.79 (d, *J* = 47.6 Hz, 2H), 3.59 (br s, 2H), 2.62-2.42 (m, 2H), 2.02-1.74 (m, 6H), 1.69-0.92 (m, 26H), 0.68 (s, 3H); $^{13}$C NMR (100 MHz, $CDCl_3$) δ 207.59 (d, *J* = 19.2 Hz), 84.92 (d, *J* = 184.2), 71.42 (d, *J* = 8.0 Hz), 71.30, 55.71, 54.87,

43.79, 43.76, 42.44, 40.12, 39.17, 37.32, 36.87, 35.22, 35.17, 34.92, 34.07, 30.35, 28.70, 28.58, 26.86, 23.36, 21.16, 18.50, 12.13, 12.11; HRMS (*m/z*): $[M + HCOO - H]^-$ calcd. for $C_{26}H_{42}FO_5$, 453.3022; found, 453.3025.

**AAA-7 (11)**. Synthesis and characterization previously described. [3]

**AAA-9 (7).** 38 mg (52% over 2 steps); TLC $R_f$ = 0.45 (40% ethyl acetate/60% hexanes); $^1$H NMR (400 MHz, $CDCl_3$) δ 4.77 (d, *J* = 48.0 Hz, 2H), 3.64-3.56 (m, 1H), 2.60-2.39 (m, 2H), 1.95-1.48 (m, 9H), 1.41-0.89 (m, 24H), 0.62 (s, 3H); $^{13}$C NMR (100 MHz, $CDCl_3$) δ 207.62 (d, *J* = 19.1 Hz), 84.89 (d, *J* = 184.2), 71.79 (d, *J* = 6.4 Hz), 56.47, 55.89, 42.73, 40.08, 40.42, 40.15, 36.42, 35.83, 35.34, 35.25, 35.18, 34.55, 30.51, 28.70, 28.16, 27.17, 26.40, 24.17, 23.34, 20.80, 18.37, 12.03, 12.01; HRMS (*m/z*): $[M + HCOO - H]^-$ calcd. for $C_{26}H_{42}FO_4$, 437.3073; found, 437.3088.

**General procedure for the sulfonation to provide the gut-restricted α-fluoromethyl ketone analogs.**

To a 0.1 M solution of the alcoholic precursor (1.0 equiv.) in pyridine, $SO_3$.pyridine (2.0 equiv.) was added and the resulting solution was stirred at rt for 18 h. The reaction mixture was concentrated using a rotary evaporator. The resulting slurry was resuspended in 5 mL of 10:1 dichloromethane:methanol and 5 mL of a saturated solution of sodium bicarbonate was added. The biphasic solution was concentrated using a rotary evaporator. The resulting crude compound was then purified by silica gel chromatography (80% dichloromethane/20% methanol) to provide the sulfonated compound.

**AAA-2’** (**3**). Yield = 12 mg (20% di-sulfonated) along with 40 mg (50%) of **AAA-2** was also obtained; TLC $R_f$ = 0.15 (20% methanol/80% dichloromethane); $^1$H NMR (400 MHz, $CD_3OD$) δ 4.90 (d, *J* = 47.6 Hz, 2H), 4.47 (d, *J* = 2.4 Hz, 1H), 4.21-4.14 (m, 1H), 2.57-2.26 (m, 4H), 2.15-1.72 (m, 9H), 1.67-1.02 (m, 13H), 0.95-0.94 (m, 6H), 0.69 (S, 3H); $^{13}$C NMR (100 MHz, $CD_3OD$) δ 207.69 (d, *J* = 17.0 Hz), 84.51 (d, *J* = 180.9), 79.60, 77.22, 55.71, 49.85, 42.36, 41.72, 39.39, 39.20, 35.54, 35.20, 34.88, 34.39, 34.00, 33.27, 30.27, 28.59, 27.71, 27.62, 22.92, 21.77, 20.32, 17.52, 10.77; HRMS (*m/z*): $[M - H]^-$ calcd. for $C_{25}H_{40}FO_9S_2$, 567.2103; found, 567.2114. Also seen $[M – H/2]^-$ calcd. for $C_{25}H_{39}FO_9S_2$, 283.1015; found, 283.1019.

**AAA-2** (GR-7; **2**). Synthesis and characterization previously described. [3]

**AAA-4** (**4**). Yield = 25 mg (70% brsm); TLC $R_f$ = 0.54 (20% methanol/80% dichloromethane); $^1$H NMR (400 MHz, $CD_3OD$) δ 4.91 (d, *J* = 47.6 Hz, 2H), 4.32-4.24 (m, 1H), 3.94 (s, 1H), 2.59-2.39 (m, 2H), 1.99-1.72 (m, 9H), 1.66-1.27 (m, 13H), 1.18-0.98 (m, 6H), 0.93 (s, 3H), 0.70 (s, 3H); $^{13}$C NMR (100 MHz, $CD_3OD$) δ 207.77 (d, *J* = 16.8 Hz), 84.51 (d, *J* = 181.1), 79.22, 72.57, 47.83, 46.59, 46.13, 42.32, 36.00, 35.20, 34.94, 34.04, 33.79, 33.33, 33.17, 28.54, 28.47, 27.29, 27.17, 26.87, 25.97, 23.42, 22.16, 16.27, 11.76; HRMS (*m/z*): $[M - H]^-$ calcd. for $C_{25}H_{40}FO_6S$, 487.2535; found, 487.2541.

**AAA-6** (**5**). Yield = 13 mg (40% brsm); TLC $R_f$ = 0.36 (20% methanol/80% dichloromethane); $^1$H NMR (400 MHz, $CD_3OD$) δ 4.91 (d, *J* = 47.2 Hz, 2H), 4.27-4.19 (m, 1H), 3.51-3.44 (m, 1H), 2.58-2.38 (m, 2H), 2.04-2.01 (m, 1H), 1.92-1.68 (m, 8H), 1.58-1.03

(m, 16H), 0.97-0.94 (m, 6H), 0.71 (s, 3H); $^{13}$C NMR (100 MHz, $CD_3OD$) δ 207.72 (d, *J* = 17.0 Hz), 84.51 (d, *J* = 181.1 Hz), 78.56, 70.39, 55.99, 55.01, 43.36, 43.04, 42.71, 40.08, 39.23, 37.06, 35.20, 34.58, 34.09, 33.99, 33.67, 28.62, 28.17, 27.29, 26.50, 22.39, 20.98, 17.61, 11.21; HRMS (*m/z*): $[M - H]^-$ calcd. for $C_{25}H_{40}FO_6S$, 487.2535; found, 487.2553.

**AAA-8** (**6**)**.** Yield = 12 mg (40% brsm); TLC $R_f$ = 0.41 (20% methanol/80% dichloromethane); $^1$H NMR (400 MHz, $CD_3OD$) δ 4.91 (d, *J* = 47.2 Hz, 2H), 4.18-4.10 (m, 1H), 3.93 (s, 1H), 3.79 (d, *J* = 2.8 Hz, 1H), 2.59-2.39 (m, 3H), 2.31-2.24 (m, 1H), 2.04-1.70 (m, 9H), 1.64-1.51 (m, 5H), 1.44-1.28 (m, 6H), 1.16-0.99 (m, 5H), 0.92 (s, 3H), 0.71 (s, 3H); $^{13}$C NMR (100 MHz, $CD_3OD$) δ 207.81 (d, *J* = 15.3 Hz), 84.52 (d, *J* = 180.8 Hz), 79.49, 72.60, 67.54, 46.53, 46.04, 41.88, 41.57, 39.54, 36.29, 35.25, 34.98, 34.39, 34.34, 34.04, 28.57, 28.17, 27.45, 27.19, 26.43, 22.79, 21.63, 16.33, 11.54; HRMS (*m/z*): $[M - H]^-$ calcd. for $C_{25}H_{40}FO_7S$, 503.2484; found, 503.2486.

**AAA-10** (**8**)**.** Yield = 0.65 g (34%); TLC $R_f$ = 0.12 (10% methanol/90% dichloromethane); $^1$H NMR (400 MHz, DMSO-$d_6$) δ 5.00 (d, *J* = 47.2 Hz, 2H), 3.99-3.91 (m, 1H), 2.48-2.26 (m, 2H), 1.90-1.51 (m, 9H), 1.32-0.97 (m, 16H), 0.92-0.83 (m, 7H), 0.59 (s, 3H); $^{13}$C NMR (100 MHz, DMSO-$d_6$) δ 206.55 (d, *J* = 15.3 Hz), 85.21 (d, *J* = 178.5 Hz), 76.01, 56.43, 55.81, 42.72, 42.04, 40.34, 40.08, 35.81, 35.50, 35.11, 34.56, 34.20, 33.79, 28.74, 28.10, 27.25, 26.52, 24.28, 23.63, 20.84, 18.73, 12.33, 12.31; HRMS (*m/z*): $[M - H]^-$ calcd. for $C_{25}H_{40}FO_5S$, 471.2580; found, 471.2591.

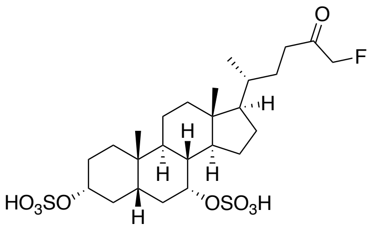
O
F
H
H
H
H
H
HO3SO
OSO3H

**3**

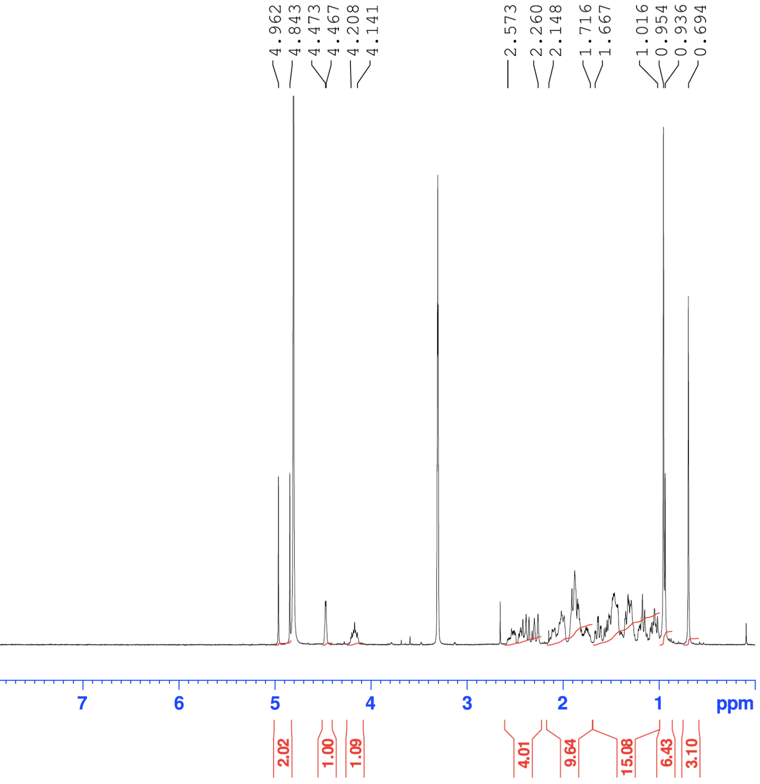
4.962
4.843
4.473
4.467
4.208
4.141
2.573
2.260
2.148
1.716
1.667
1.016
0.954
0.936
0.694
11
10
9
8
7
6
5
4
3
2
1
ppm
2.02
1.00
1.09
4.01
9.64
15.08
6.43
3.10

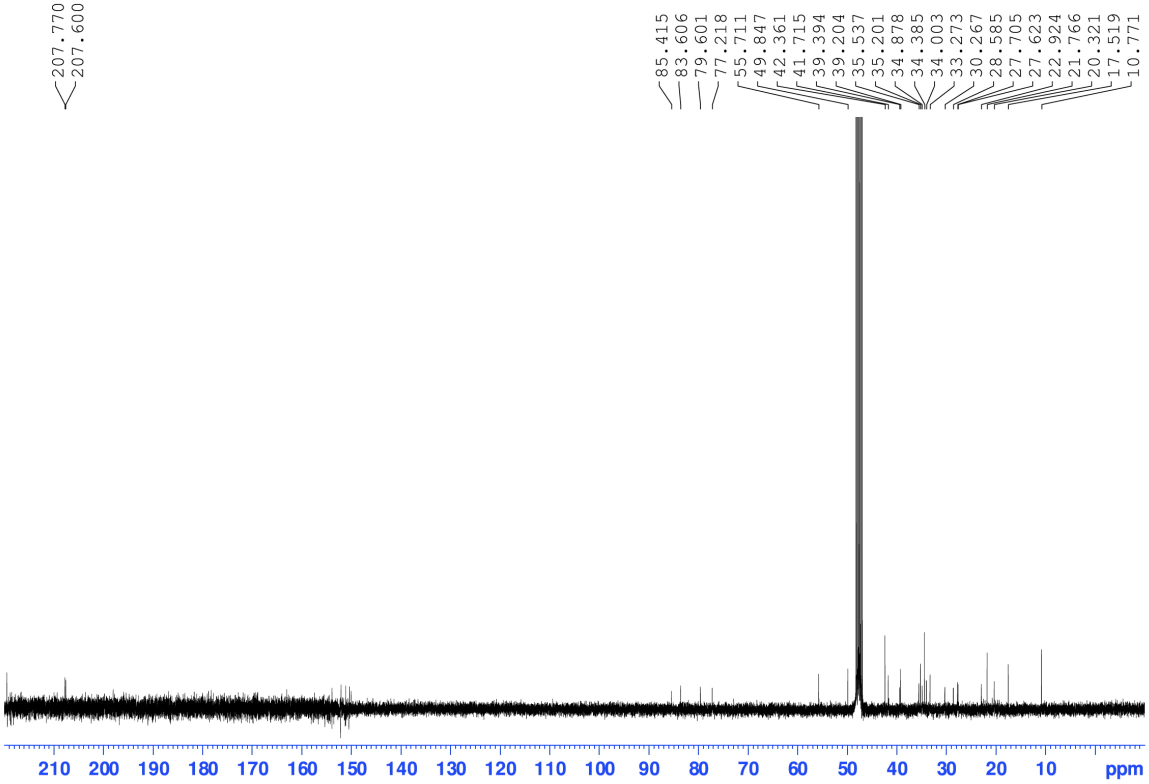
207.770
207.600
85.415
83.606
79.601
77.218
55.711
49.847
42.361
41.715
39.394
39.204
35.537
35.201
34.878
34.385
34.003
33.273
30.267
28.585
27.705
27.623
22.924
21.766
20.321
17.519
10.771
210
200
190
180
170
160
150
140
130
120
110
100
90
80
70
60
50
40
30
20
10
ppm

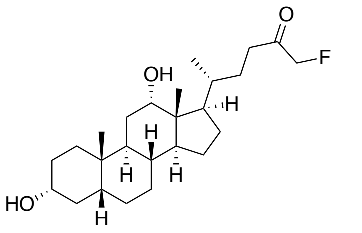
O
F
OH
H
HO

**9**

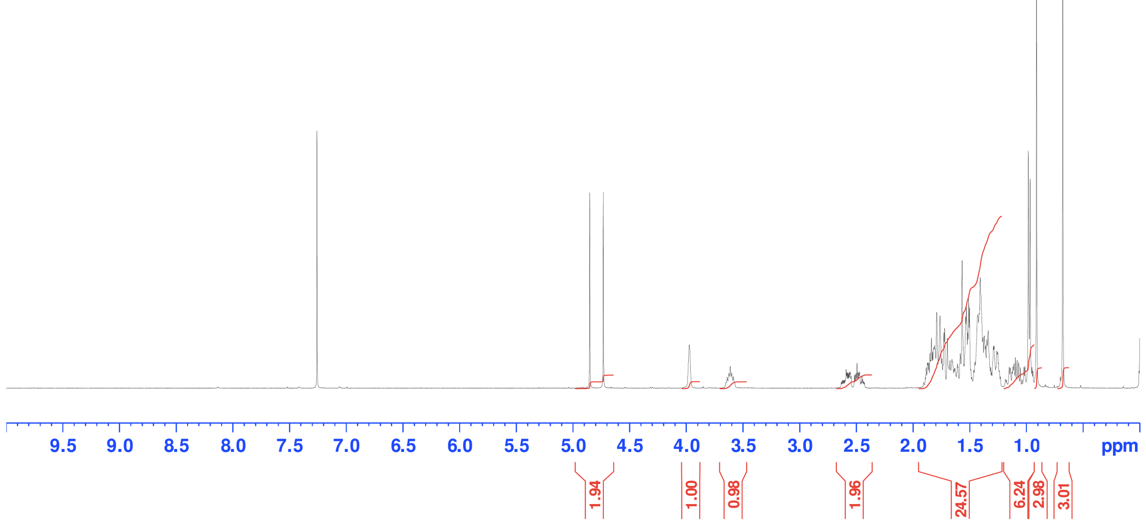
4.851
4.732
3.972
3.650
3.573
2.632
2.427
1.905
1.228
1.184
0.941
0.909
0.678
9.5 9.0 8.5 8.0 7.5 7.0 6.5 6.0 5.5 5.0 4.5 4.0 3.5 3.0 2.5 2.0 1.5 1.0 ppm
1.94
1.00
0.98
1.96
24.57
6.24
2.98
3.01

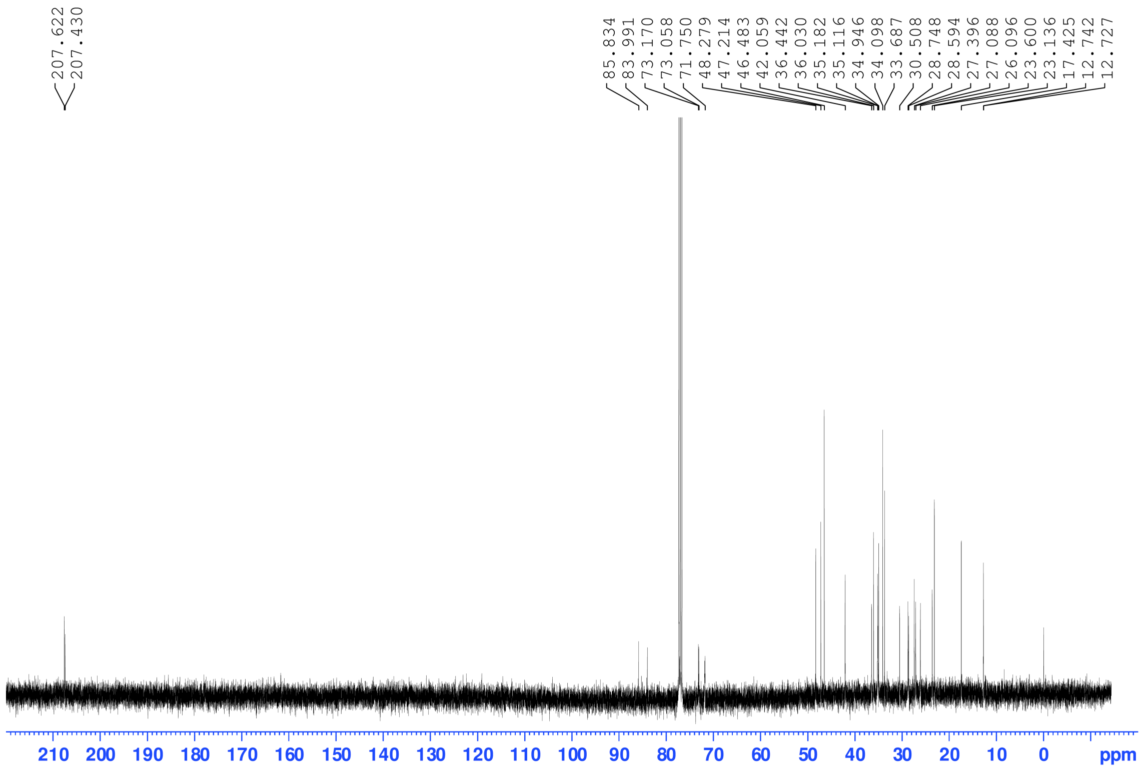
207.622
207.430
85.834
83.991
73.170
73.058
71.750
48.279
47.214
46.483
42.059
36.442
36.030
35.182
35.116
34.946
34.098
33.687
30.508
28.748
28.594
27.396
27.088
26.096
23.600
23.136
17.425
12.742
12.727
210 200 190 180 170 160 150 140 130 120 110 100 90 80 70 60 50 40 30 20 10 0 ppm

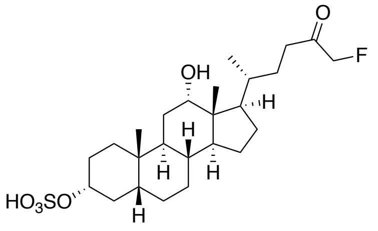
O
F
OH
H
$HO_3SO$

**4**

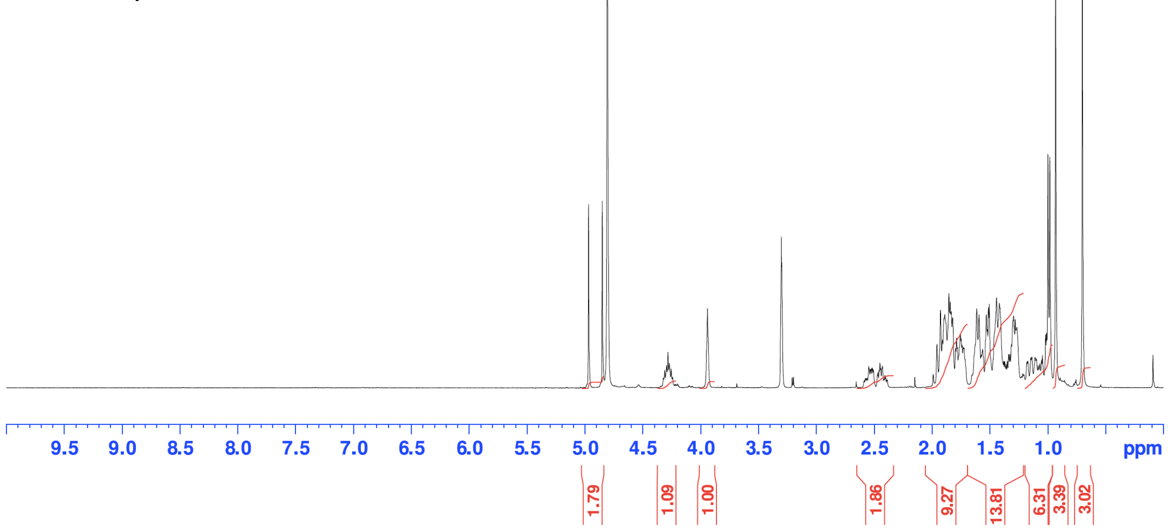
4.969
4.850
4.323
4.243
3.942
2.585
2.386
1.990
1.724
1.655
1.267
1.181
0.983
0.932
0.701
9.5 9.0 8.5 8.0 7.5 7.0 6.5 6.0 5.5 5.0 4.5 4.0 3.5 3.0 2.5 2.0 1.5 1.0 ppm
1.79
1.09
1.00
1.86
9.27
13.81
6.31
3.39
3.02

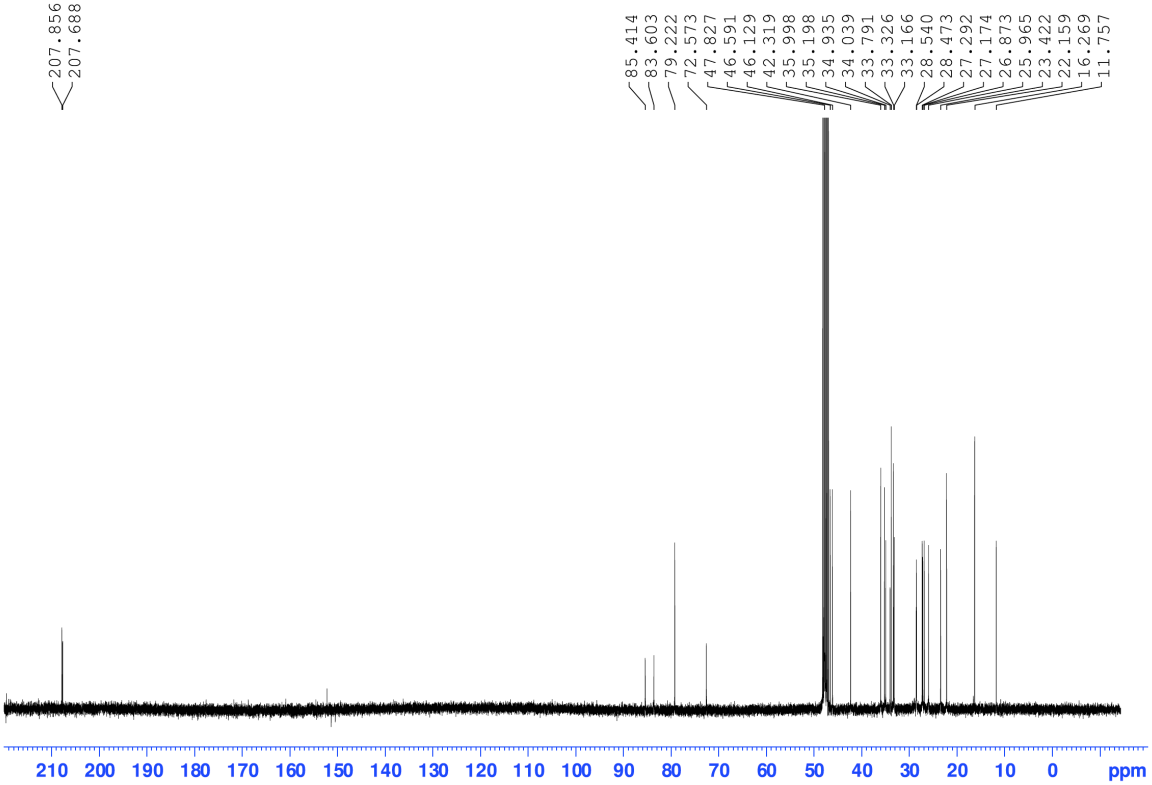
207.856
207.688
85.414
83.603
79.222
72.573
47.827
46.591
46.129
42.319
35.998
35.198
34.935
34.039
33.791
33.326
33.166
28.540
28.473
27.292
27.174
26.873
25.965
23.422
22.159
16.269
11.757
210 200 190 180 170 160 150 140 130 120 110 100 90 80 70 60 50 40 30 20 10 0 ppm

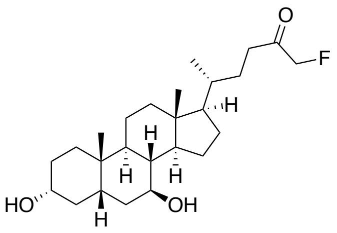
O
F
H
HO
OH
H

**10**

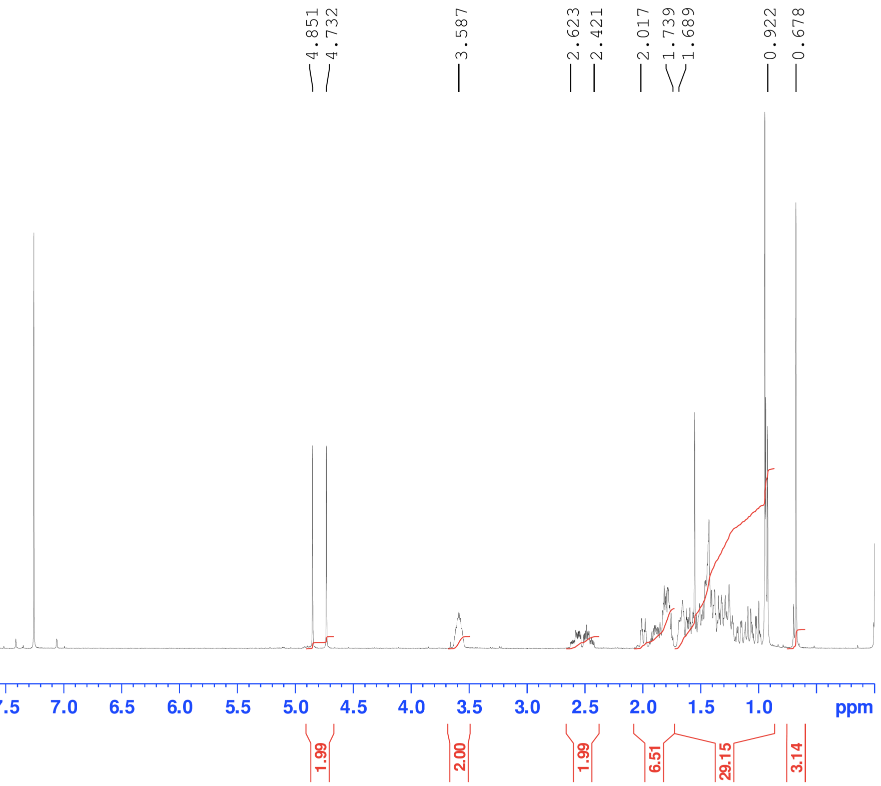
4.851
4.732
3.587
2.623
2.421
2.017
1.739
1.689
0.922
0.678
9.5 9.0 8.5 8.0 7.5 7.0 6.5 6.0 5.5 5.0 4.5 4.0 3.5 3.0 2.5 2.0 1.5 1.0 ppm
1.99
2.00
1.99
6.51
29.15
3.14

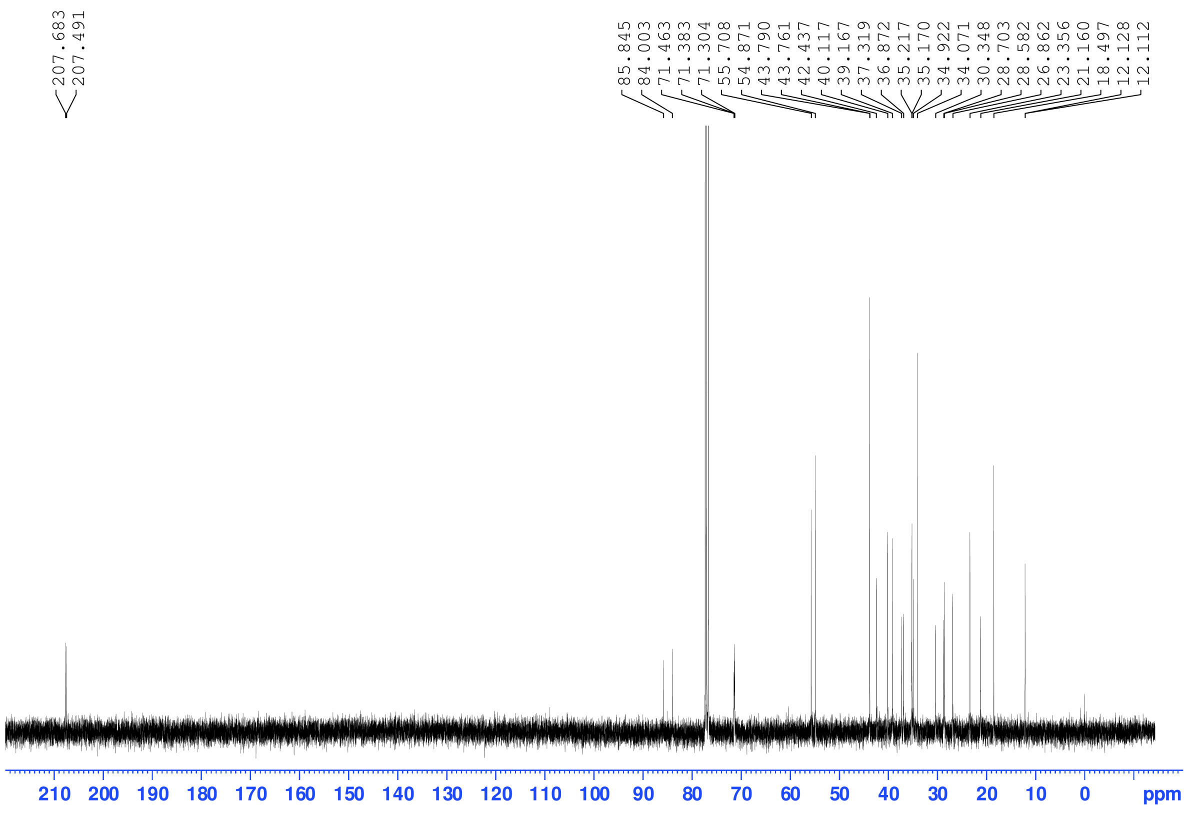
207.683
207.491
85.845
84.003
71.463
71.383
71.304
55.708
54.871
43.790
43.761
42.437
40.117
39.167
37.319
36.872
35.217
35.170
34.922
34.071
30.348
28.703
28.582
26.862
23.356
21.160
18.497
12.128
12.112
210 200 190 180 170 160 150 140 130 120 110 100 90 80 70 60 50 40 30 20 10 0 ppm

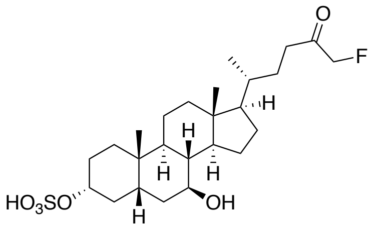
O
F
H
HO3SO
OH

**5**

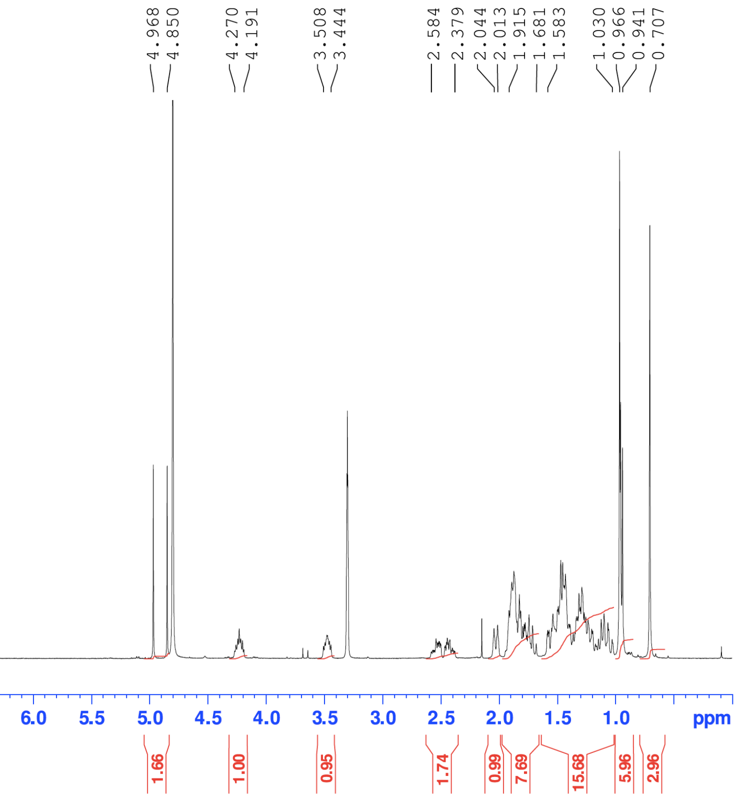
4.968
4.850
4.270
4.191
3.508
3.444
2.584
2.379
2.044
2.013
1.915
1.681
1.583
1.030
0.966
0.941
0.707
9.5
9.0
8.5
8.0
7.5
7.0
6.5
6.0
5.5
5.0
4.5
4.0
3.5
3.0
2.5
2.0
1.5
1.0
ppm
1.66
1.00
0.95
1.74
0.99
7.69
15.68
5.96
2.96

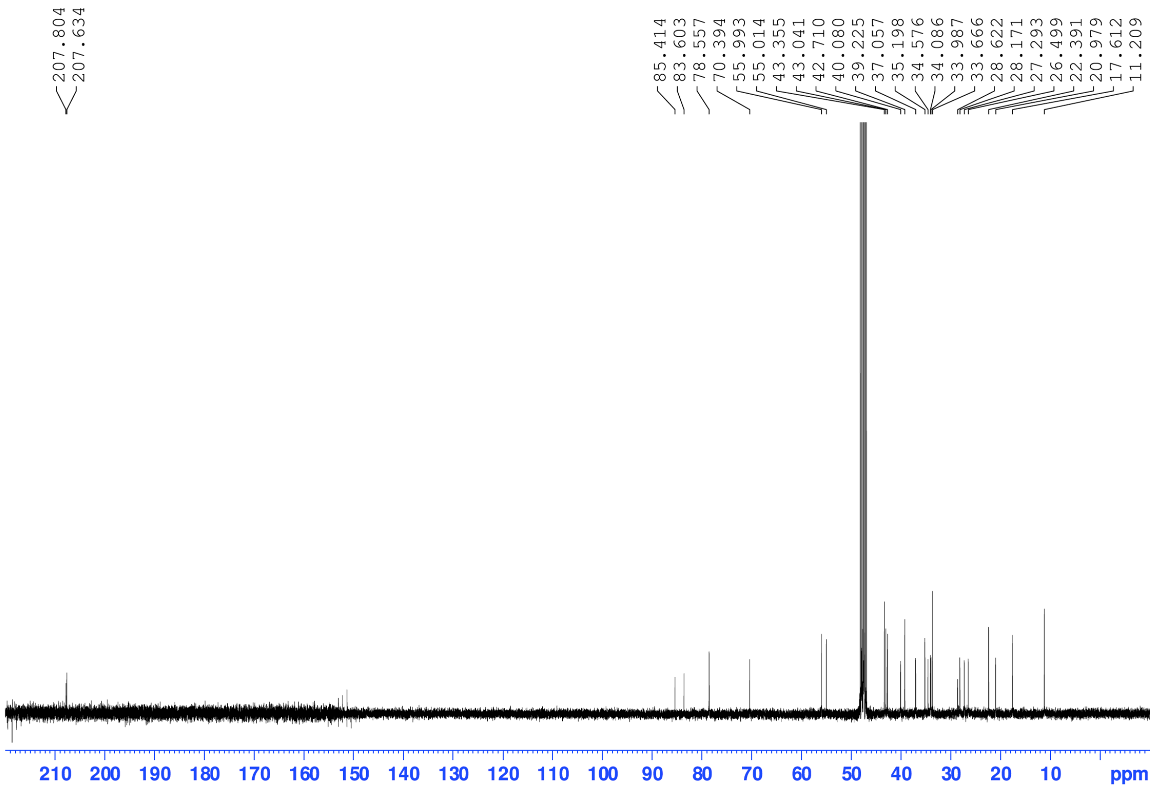
207.804
207.634
85.414
83.603
78.557
70.394
55.993
55.014
43.355
43.041
42.710
40.080
39.225
37.057
35.198
34.576
34.086
33.987
33.666
28.622
28.171
27.293
26.499
22.391
20.979
17.612
11.209
210
200
190
180
170
160
150
140
130
120
110
100
90
80
70
60
50
40
30
20
10
ppm

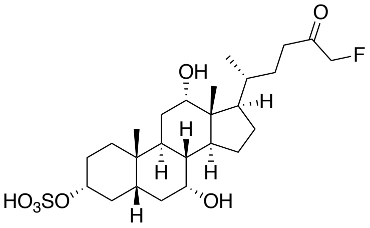
O
F
OH
H
H
H
H
HO3SO
OH
H

**6**

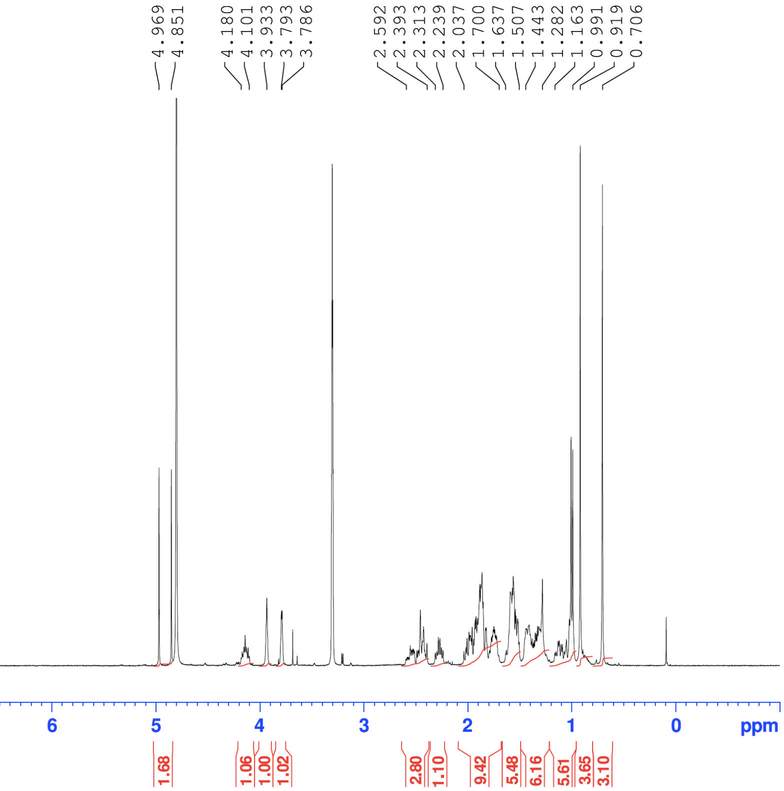
4.969
4.851
4.180
4.101
3.933
3.793
3.786
2.592
2.393
2.313
2.239
2.037
1.700
1.637
1.507
1.443
1.282
1.163
0.991
0.919
0.706
9
8
7
6
5
4
3
2
1
0
ppm
1.68
1.06
1.00
1.02
2.80
1.10
9.42
5.48
6.16
5.61
3.65
3.10

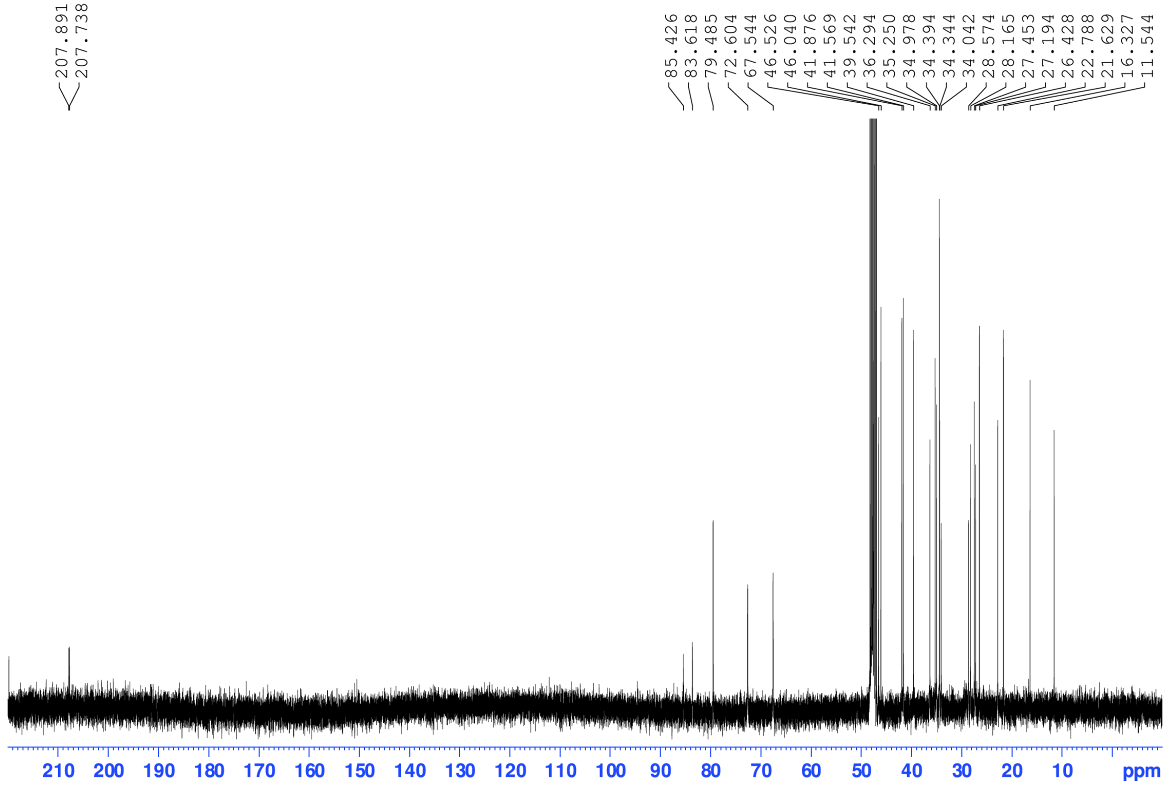
207.891
207.738
85.426
83.618
79.485
72.604
67.544
46.526
46.040
41.876
41.569
39.542
36.294
35.250
34.978
34.394
34.344
34.042
28.574
28.165
27.453
27.194
26.428
22.788
21.629
16.327
11.544
210
200
190
180
170
160
150
140
130
120
110
100
90
80
70
60
50
40
30
20
10
ppm

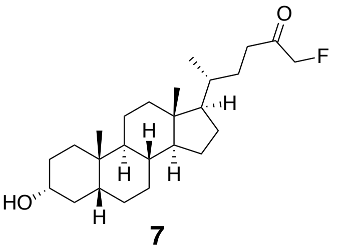
O
F
H
H
H
H
HO
H

**7**

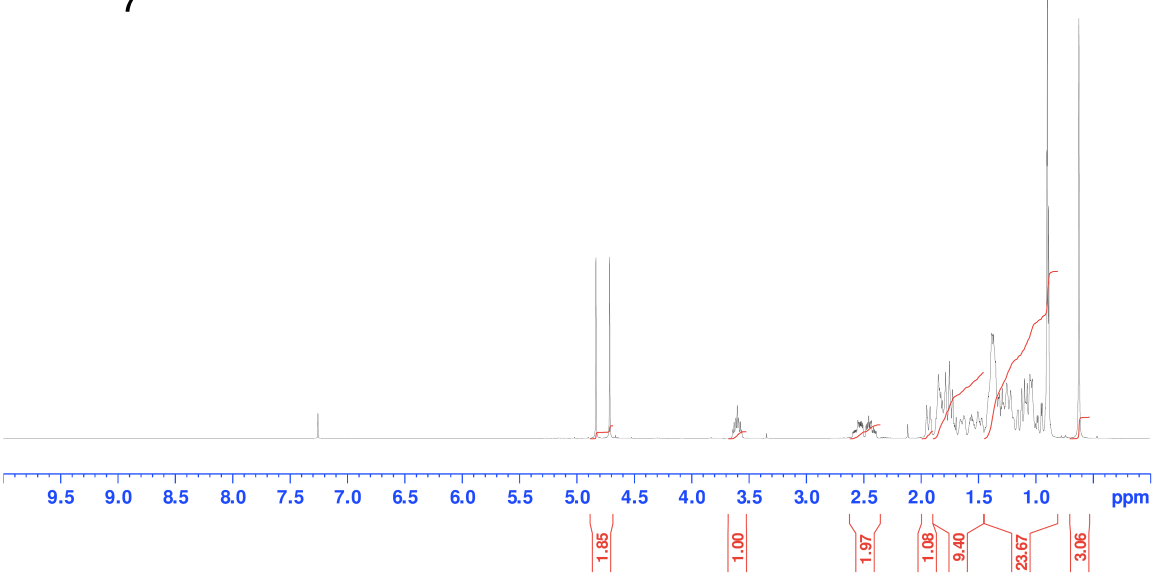
4.834
4.714
3.641
3.563
2.595
2.391
1.951
1.475
1.414
0.888
0.624
9.5 9.0 8.5 8.0 7.5 7.0 6.5 6.0 5.5 5.0 4.5 4.0 3.5 3.0 2.5 2.0 1.5 1.0 ppm
1.85
1.00
1.97
1.08
9.40
23.67
3.06

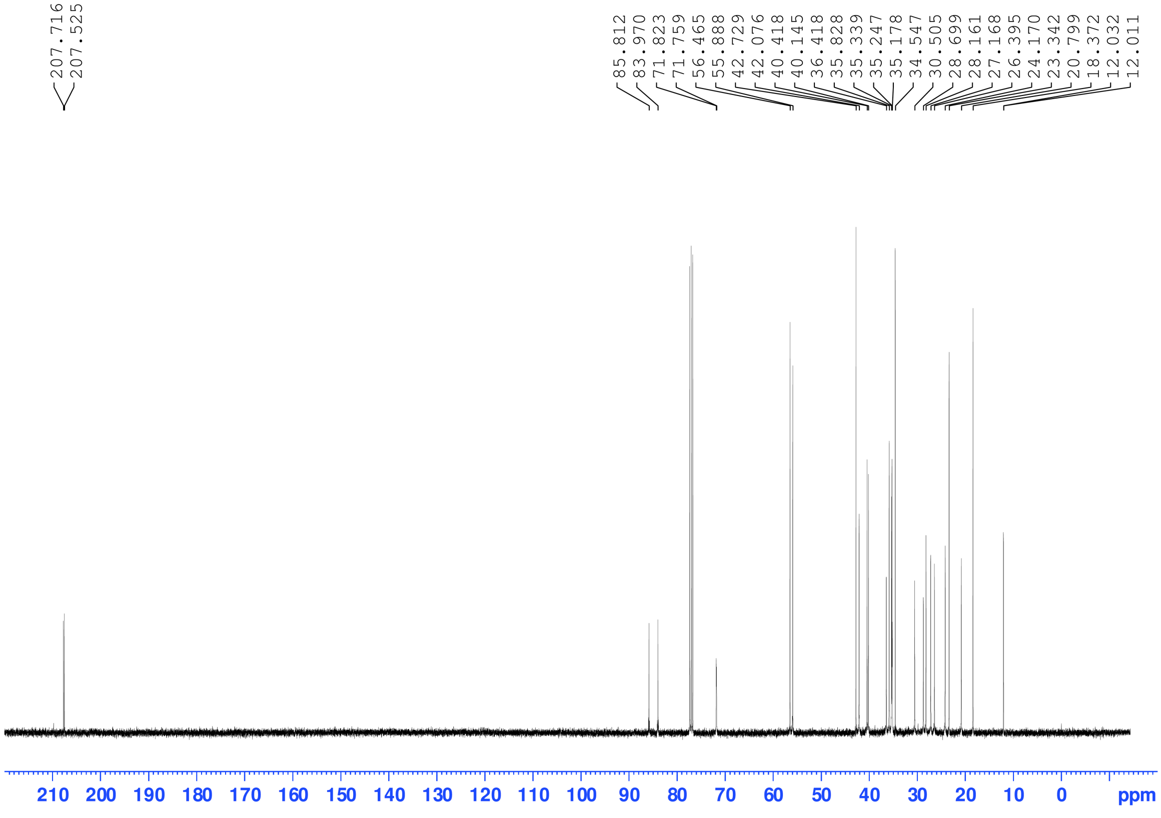
207.716
207.525
85.812
83.970
71.823
71.759
56.465
55.888
42.729
42.076
40.418
40.145
36.418
35.828
35.339
35.247
35.178
34.547
30.505
28.699
28.161
27.168
26.395
24.170
23.342
20.799
18.372
12.032
12.011
210 200 190 180 170 160 150 140 130 120 110 100 90 80 70 60 50 40 30 20 10 0 ppm

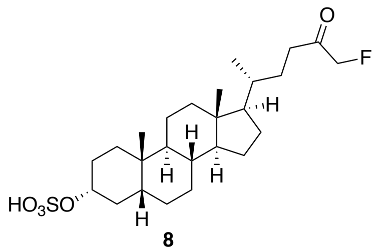
O
F
H
H
H
H
HO3SO
H

**8**
**AAA-10**

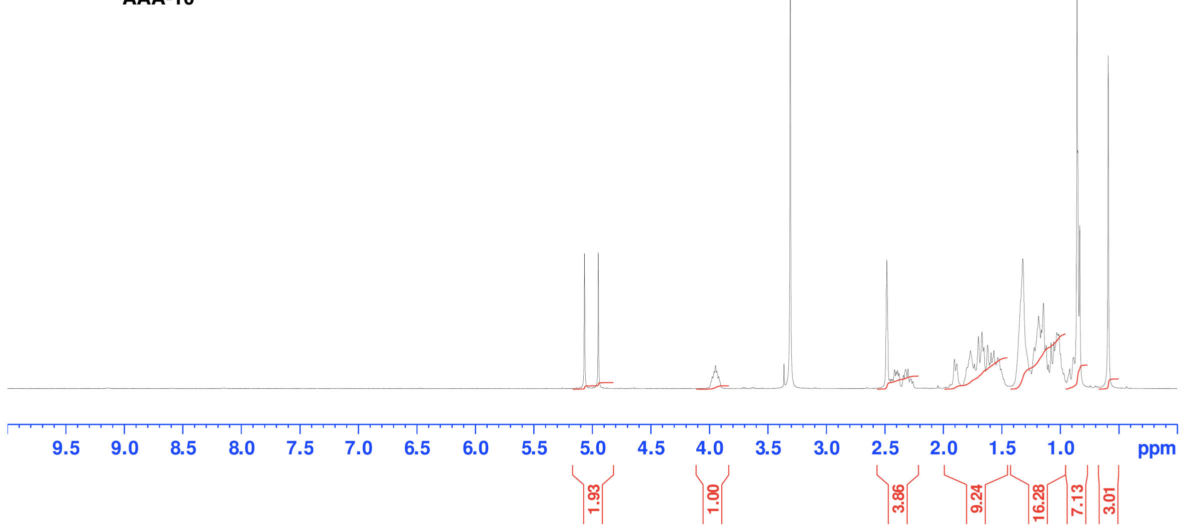
5.067
4.949
3.986
3.908
2.482
2.260
1.904
1.506
1.320
0.968
0.919
0.833
0.589
9.5 9.0 8.5 8.0 7.5 7.0 6.5 6.0 5.5 5.0 4.5 4.0 3.5 3.0 2.5 2.0 1.5 1.0 ppm
1.93
1.00
3.86
9.24
16.28
7.13
3.01

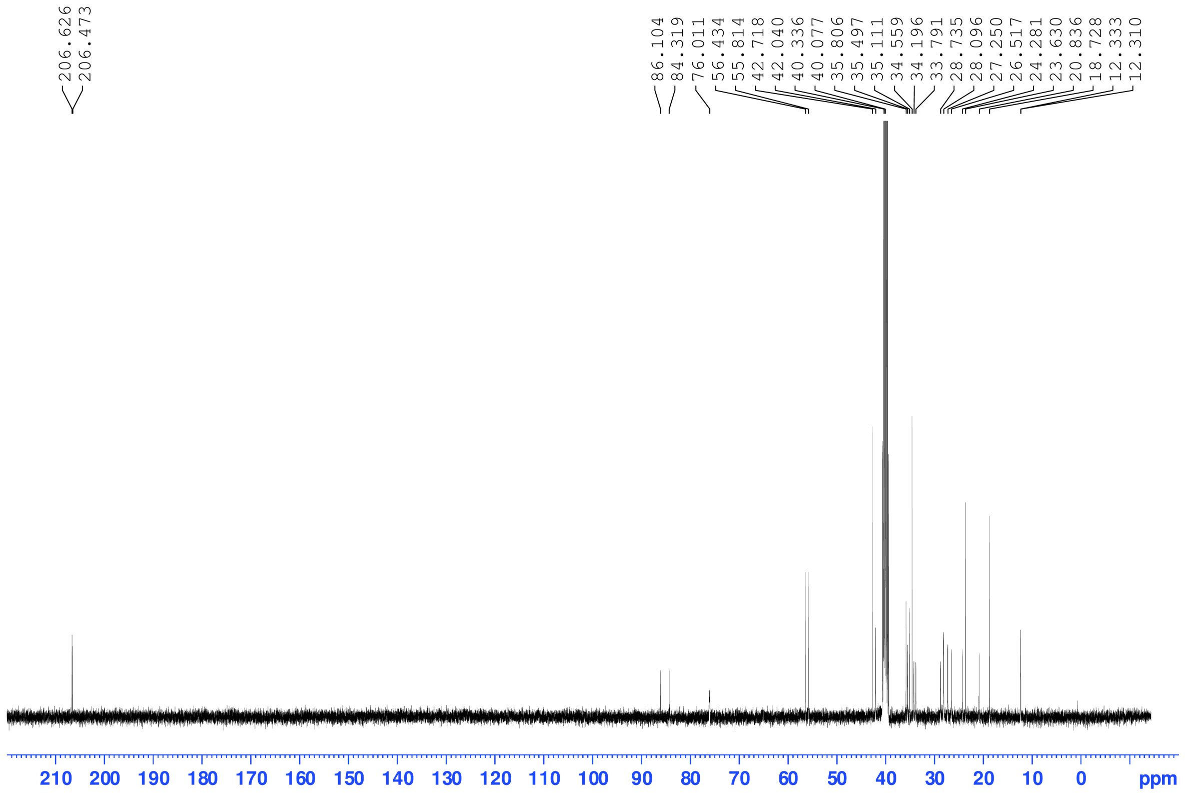
206.626
206.473
86.104
84.319
76.011
56.434
55.814
42.718
42.040
40.336
40.077
35.806
35.497
35.111
34.559
34.196
33.791
28.735
28.096
27.250
26.517
24.281
23.630
20.836
18.728
12.333
12.310
210 200 190 180 170 160 150 140 130 120 110 100 90 80 70 60 50 40 30 20 10 0 ppm

**Supporting Tables**

**Table S1. Comparison of $IC_{50}$ values against recombinant BSHs**

| Inhibitor | *B. theta* rBSH | *B. longum* rBSH |
|---|---|---|
| Compound **7** **(AAA-1)** | 427 nM | 108 nM |
| GR-**7** **(AAA-2)** | 2638 nM | 1265 nM |
| **AAA-10** | 10 nM | 80 nM |

### Supporting Figures

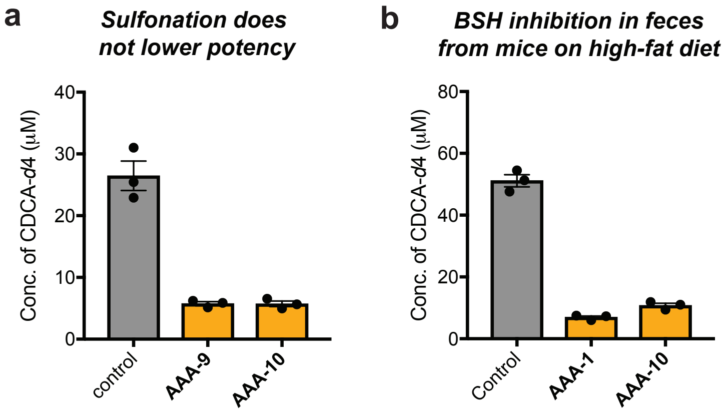

**Figure S1. Comparing BSH inhibitor efficacy in conventional mouse fecal slurry. a, AAA-10** did not lose potency due to the presence of the sulfonate group. Comparison of inhibitory activity of **AAA-10** and unsulfonated inhibitor **AAA-9** showed no difference in the inhibitory ability. Inhibitors were tested at a concentration of 10 µM. **b,** Screen in feces obtained from conventional mice fed a high-fat diet showed that **AAA-10** inhibited BSH activity with similar potency compared to the first-generation unsulfonated inhibitor, **AAA-1**. Inhibitors were tested at a concentration of 20 µM. All assays were performed in biological triplicate, and data are presented as mean ± s.e.m.

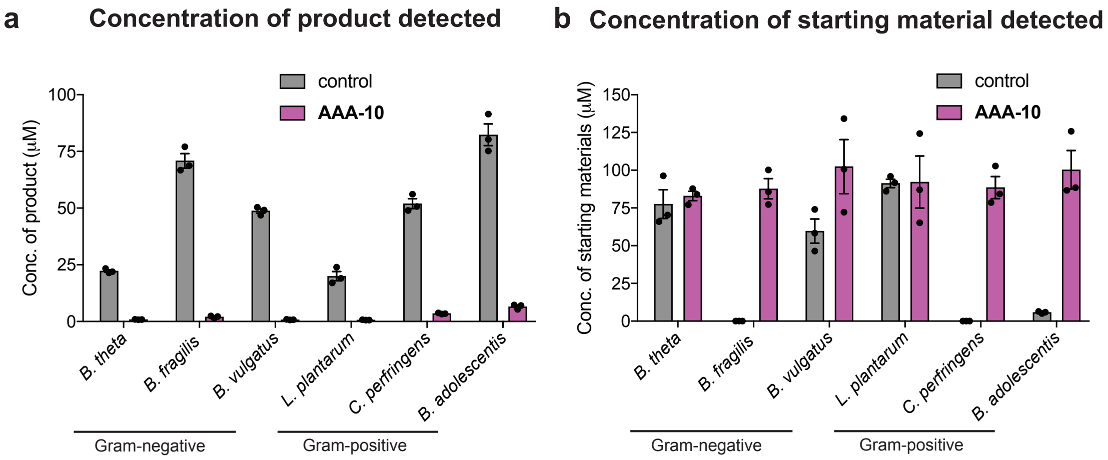

**Figure S2. Absolute bile acid concentrations for determining % deconjugation in bacterial culture assays. a,** Concentration of products formed (deconjugated bile acids) and **b,** unreacted starting materials in each culture were determined using UPLC-MS. Percent deconjugation for each sample was then determined using the following equation:

% deconjugation = Concentration of products / (Concentration of products + Concentration of starting materials) * 100.

Note that for 5 of the 6 bacteria tested, no bile acids were detected in the cultures other than the starting materials (TCA, TβMCA, TUDCA and TDCA) and their deconjugated products (i.e., CA, βMCA, UDCA and DCA). For *B. fragilis*, these bile acids and one additional compound, 7-oxo-cholic acid (7-oxo-CA), were detected. 7-oxo-CA was quantified and included in the concentration of products in (**a**). Assays were performed in biological triplicate, and all data are presented as mean ± s.e.m.

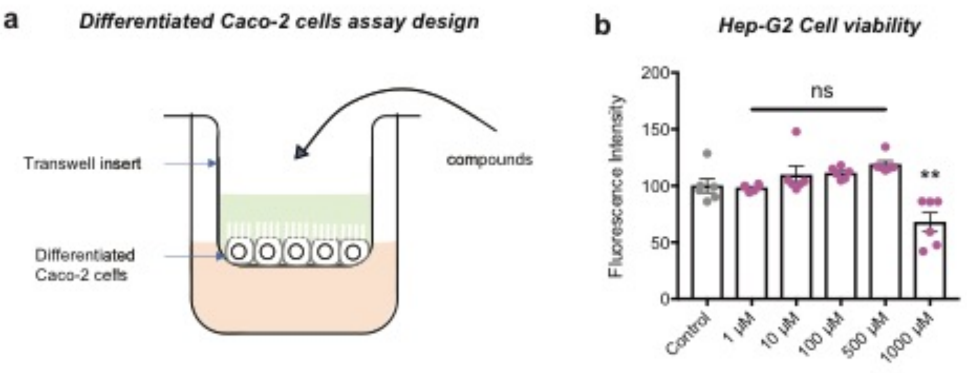

**Figure S3. Transwell assay design and AAA-10 toxicity. a,** Caco-2 cells were differentiated to form a monolayer with tight junctions. Compounds were then added to the apical side and their transport to the basolateral side was quantified by UPLC-MS. **b,** Incubation of Hep-G2 cells with **AAA-10** did not result in toxicity up to 500 μM. For **b,** one-way ANOVA followed by Dunnett's multiple comparisons test was performed, n=6, and data are presented as mean ± s.e.m.

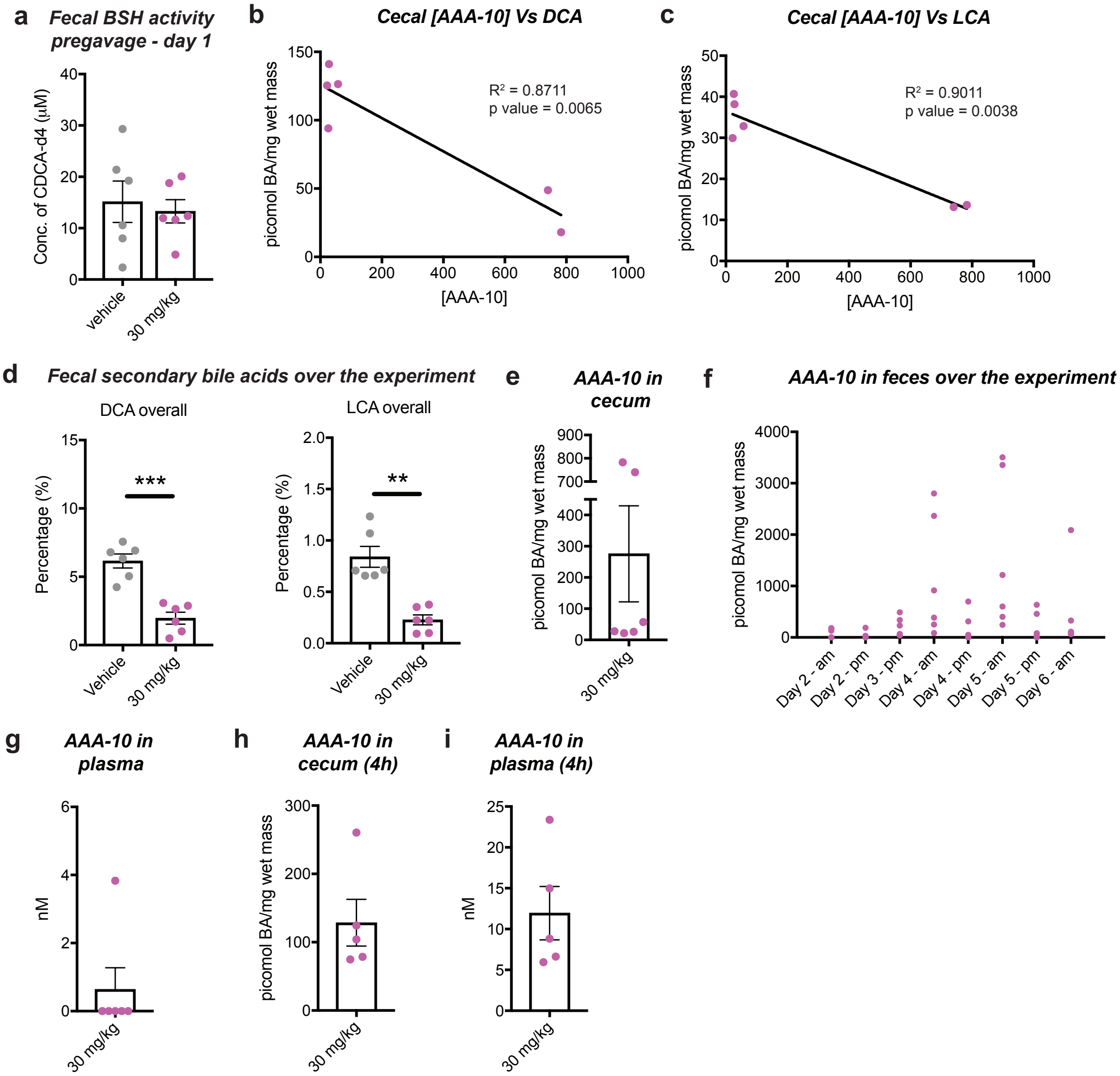

**Figure S4. AAA-10 inhibited BSH activity and reduced of secondary bile acid abundance in vivo. a,** There was no difference in BSH activity in feces pre-**AAA-10** treatment. **b** and **c,** Cecal **AAA-10** concentration was inversely correlated with the cecal concentrations of the secondary bile acids DCA and LCA. Each dot represents the concentration of **AAA-10** vs bile acid in each mouse. **d,** The abundances of DCA and LCA in feces collected over the entire period of the experiment were significantly lower in AAA-10-treated mice compared to control-treated mice. Each dot represents a mouse. See methods section for the equation used to calculate these overall abundances. **e**, Quantification of **AAA-10** in mouse cecal contents. **AAA-10** was detected in cecum 16 hours after administration. **f, AAA-10** concentration in feces collected over the period of study. Each dot represents the amount of **AAA-10** in each mouse at a particular timepoint. **g**, Quantification of **AAA-10** in mouse plasma 16h after final gavage. **AAA-10** was detected in the plasma of only 1 mouse indicating that **AAA-10** exhibits minimal systemic exposure. **h** and **i,** Quantification of **AAA-10** in a separate mouse experiment 4h post-gavage. While substantial amounts of **AAA-10** were detected in mouse cecal contents, trace amounts of **AAA-10** were detected in mouse plasma. For **a-g,** n=6 mice/group. For **b** and **c,** linear regression was performed to determine the $R^2$ and p value. For **h** and **i,** n=5 mice/group. For **a** and **d,** two-tailed Welch's t test was performed. *p<0.05, **p<0.01, ***p<0.001, ****p<0.0001, ns = not significant. All data are presented as mean ± s.e.m.

### Supporting Schemes

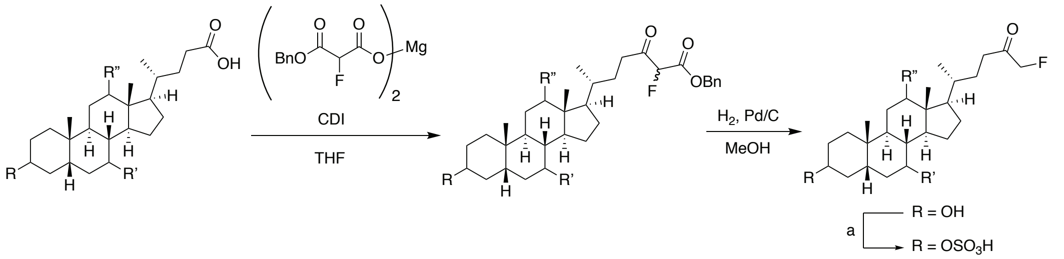

**Scheme 1: Generic scheme for synthesis of compounds 1-5.** CDI = carbonyldiimidazole, a = $SO_3$.pyridine, pyridine

**Scheme S1**. Generic scheme for the synthesis of compounds 3-8. CDI – carbonyldiimidazole, a=SO3.pyridine, pyridine.

### Supporting References

1. Yao, L. *et al*. A selective gut bacterial bile salt hydrolase alters host metabolism. *eLife* **7,** 675 (2018).
2. Xie, C. *et al*. An Intestinal Farnesoid X Receptor-Ceramide Signaling Axis Modulates Hepatic Gluconeogenesis in Mice. *Diabetes* **66,** 613–626 (2017).
3. Adhikari, A. A. *et al*. Development of a covalent inhibitor of gut bacterial bile salt hydrolases. *Nature Chemical Biology* **16,** 318–326 (2020).
4. Chaudhari, S. N. *et al*. A microbial metabolite remodels the gut-liver axis following bariatric surgery. *Cell Host & Microbe* **317,** 571 (2020).
5. Palmer, J. T. & Inc, P. Process for forming a fluoromethyl ketone. (1994).
